# Stress-induced phospho-ubiquitin formation causes parkin degradation

**DOI:** 10.1101/484857

**Authors:** Lyudmila Kovalchuke, Eugene V. Mosharov, Oren A. Levy, Lloyd A. Greene

## Abstract

Mutations in the E3 ubiquitin ligase parkin are the most common known cause of autosomal recessive parkinsonism. Multiple types of stress decrease parkin protein levels, an effect that may be relevant to sporadic Parkinson’s disease (PD), but the mechanism(s) involved in this loss remain largely unclear. We sought to elucidate these mechanisms using a PD-relevant stressor, L-DOPA, the precursor to dopamine, which forms reactive oxygen species (ROS) as well as toxic quinones via auto-oxidation. We find that L-DOPA causes parkin loss through both an oxidative stress-independent and an oxidative stress-dependent pathway. Characterization of the latter reveals that it requires both the kinase PINK1 and parkin’s interaction with phosphorylated ubiquitin (phospho-Ub) and is mediated by proteasomal degradation.

Surprisingly, mitochondrial parkin activity and autoubiquitination as well as mitophagy are not required for such loss. During stress induced by the oxidative stressor hydrogen peroxide or the metabolic uncoupler CCCP, parkin degradation also requires its association with phospho-Ub, indicating that this mechanism is broadly generalizable. As oxidative stress, metabolic dysfunction and phospho-Ub levels are all elevated in PD patients, we suggest that these changes may lead to the loss of parkin expression in PD.

## Introduction

Parkinson’s disease (PD) is a debilitating neurodegenerative disorder, affecting roughly 2% of those over the age of 80 [1]. Though most cases of PD appear to be “sporadic”, a minority of cases have a clear autosomal dominant or recessive inheritance pattern [2]. The most common known cause of autosomal recessive PD is homozygous inactivation of *PARK2*, which encodes the E3 ubiquitin ligase parkin [3]. Parkin has been demonstrated to promote cell survival in many different contexts [4]–[21]. There is also evidence that parkin loss may play a role in the pathogenesis of sporadic PD [21]–[23]. As such, strategies aimed at upregulating parkin levels or maintaining it in an active state hold therapeutic promise for sporadic PD [16], [24]–[26].

Though much work has examined the functions of parkin, and, more recently, the mechanisms of its activation by phospho-ubiquitin (phospho-Ub) and phosphorylation [27], substantially less is known about how levels of both total and activated parkin are regulated in cells. An important insight into parkin regulation comes from *in vitro* and *in vivo* observations that diverse stressors cause a decrease in parkin protein levels [14], [15], [28]–[31]. These stressors include mitochondrial complex I inhibitors [15], [28]–[30], oxidative agents [14], [15], [29], [30], and a DNA-damaging agent [31]. Mitochondrial dysfunction and oxidative stress are well-characterized aspects of PD [32]–[35], suggesting that parkin loss from these stresses may occur in, and possibly contribute to, the progression of this disorder. However, the mechanism(s) involved in parkin loss from these stressors are largely unclear.

Additionally, mitochondrial depolarization has also been shown to cause parkin loss. This loss is generally thought to be linked to the process of parkin-mediated mitophagy [36]–[39], though one study has suggested that parkin’s autoubiquitination leads to its degradation and prevents mitophagy following mitochondrial depolarization [40]. The degree to which parkin loss from mitochondrial depolarization aligns mechanistically with parkin loss from other stressors is uncertain. One possible contributor in common is the mitochondrial kinase PINK1, which has been implicated in parkin loss from both mitochondrial depolarization and hydrogen peroxide exposure [40], [41]. PINK1 phosphorylates ubiquitin at Ser65, and the phospho-Ub in turn binds parkin, partially activating it [42]–[44]. Phospho-Ub-bound parkin itself serves as an efficient substrate for PINK1 [45]–[47], which phosphorylates it at Ser65 in its ubiquitin-like (Ubl) domain and thereby promotes its full activation [48], [49]. A well-described function for parkin activated in this manner is to poly-ubiquitinate mitochondrial proteins (subsequently referred to as “canonical” mitochondrial parkin activity), which, in concert with PINK1-mediated phosphorylation, defines a positive feedback loop that generates mitochondrial phosphorylated poly-ubiquitin (phospho-poly-Ub) chains and initiates mitophagy [50], [51]. Mitophagy results in turnover of both mitochondrial proteins and of parkin itself [36], [37]. It is, however, unclear whether parkin loss triggered by oxidative stressors utilizes mechanisms, and, in particular, what the roles of PINK1, phospho-Ub, parkin activity, parkin auto-ubiquitination, and autophagy are in this process.

In the current study, we have explored the mechanisms of parkin loss promoted by oxidative stress. For this purpose, we primarily employed L-DOPA, the precursor to dopamine (DA). L-DOPA and DA generate reactive oxygen species (ROS) as well as toxic quinones via auto-oxidation [52], [53], and there is evidence that these stressors may contribute to PD pathogenesis [32], [54], [55]. L-DOPA is also a standard therapy for PD, and the idea has been raised that, as well as providing symptomatic relief in PD, its prolonged use could also contribute to neuronal degeneration [56], [57].

We show that L-DOPA induces parkin loss through two distinct pathways: an oxidative stress-dependent pathway and an oxidative stress-independent pathway, each accounting for about half of parkin loss. We characterize the former and show that parkin’s association with PINK1-dependent phospho-Ub is critical for parkin loss via this pathway. Furthermore, we find that parkin’s association with phospho-Ub generated by other stressors also leads to parkin degradation, suggesting that this mechanism is broadly-generalizable. Surprisingly, we find that parkin loss downstream of its association with phospho-Ub does not appear to require canonical mitochondrial parkin activity or mitophagy.

## Results

### L-DOPA causes parkin degradation

To assess the effect of L-DOPA on cellular levels of parkin, we treated neuronally differentiated PC12 cells with various concentrations of L-DOPA for 24 hours and determined relative parkin expression by Western immunoblotting (WB). PC12 cells are catecholaminergic (producing principally DA) and have been widely used to investigate catecholamine function and metabolism as well as for model studies of potential causes and treatments of PD [58], [59]. Neuronally differentiated PC12 cells also possess levels of parkin that are easily detected by WB, making them a fitting model in which to evaluate the effect of stress on endogenous parkin. Upon exposure to L-DOPA, we observed a dose-dependent loss of parkin protein that reached significance at concentrations of 100 μM and beyond (Fig. 1A). Given the robust parkin loss we observed with 200 μM L-DOPA (68.4 ± 5.2% parkin remaining with 200 μM L-DOPA compared to 0 μM L-DOPA, p = 0.01, N = 5), we chose this dose for further experiments. A time course study revealed that significant parkin loss after treatment with 200 μM L-DOPA is detectable around 6 hours post-treatment and is nearly complete by 26 hours (0 μM L-DOPA: 99.2 ± 5.2% relative parkin level at 26 hours, N = 4; 200 μM L-DOPA: 55.6 ± 3.2% relative parkin level, N = 4; p = 0.03) (Fig. 1B).

**Figure 1.**
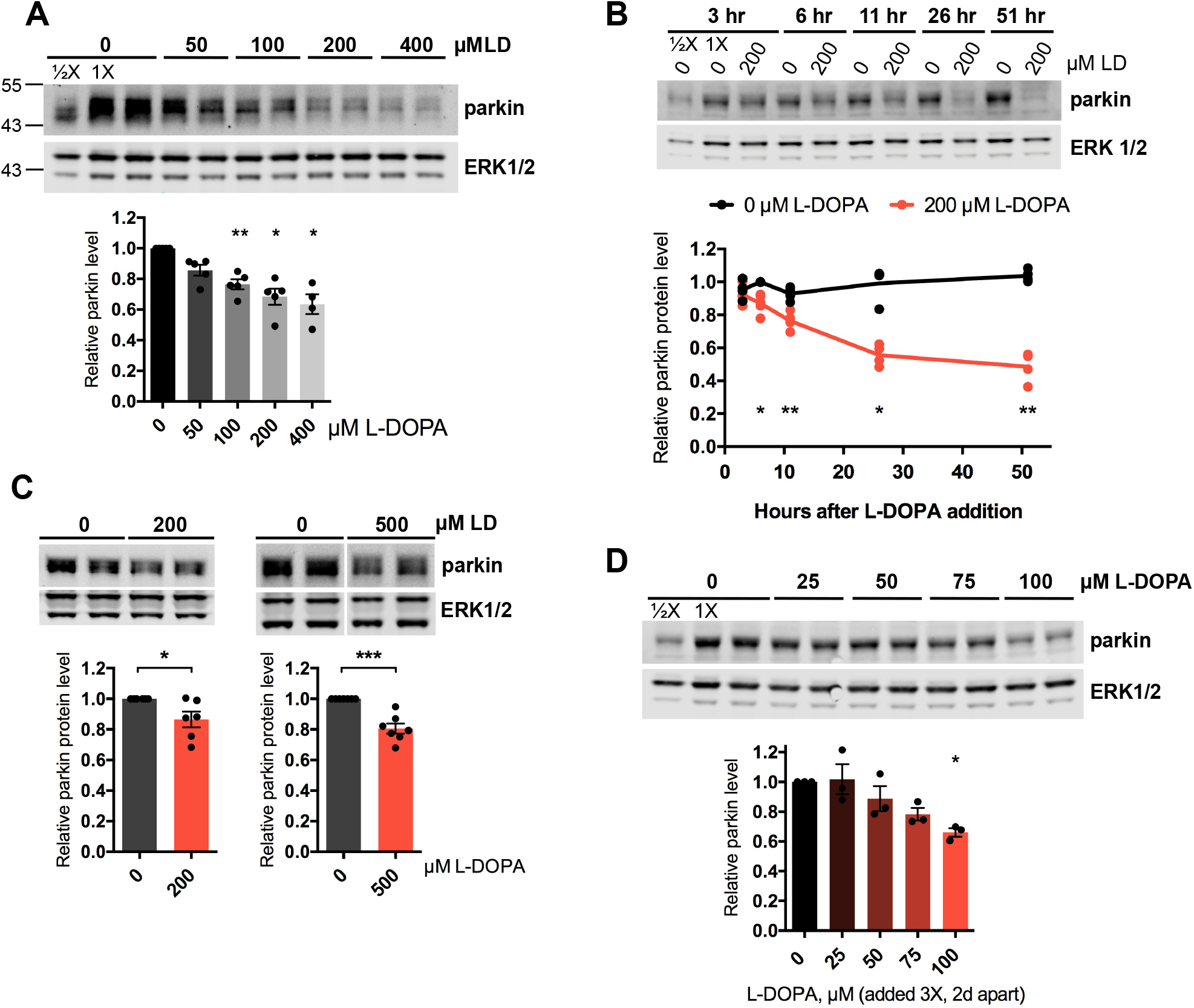
L-DOPA decreases cellular parkin protein levels. A,B. Dose-response and time-course experiments with L-DOPA in PC12 cells. Neuronally differentiated PC12 cells were treated with the indicated concentrations of L-DOPA (LD) for 24 hours (A) or with 200 μM L-DOPA for the indicated times up to 51 hours (B), and total cell lysates (in lysis buffer containing 2% LDS) were assessed for relative parkin levels by Western immunoblotting and normalization to the average of ERK 1 and 2 protein levels. Here and subsequently (though not always shown), half of the sample labelled “1X” was loaded in an adjacent well (labelled “1/2X”) to serve as a quantification standard (see methods). Representative Western immunoblots are shown here and in subsequent panels, and quantification of N = 4-5 independent experiments is shown below. C. L-DOPA decreases parkin protein levels in cultured rat cortical neurons. Neurons were treated with 200 μM L-DOPA for 24 hours (left, N = 6 independent experiments) or 500 μM L-DOPA for 15-18 hours (right, N = 7 independent experiments). D. Non-toxic concentrations of L-DOPA still induce parkin loss. Representative Western blot and quantification of parkin protein levels after exposure to the indicated concentrations of L-DOPA following the treatment paradigm described in Figure S1B. N = 3 independent experiments. A-D. Error bars show SEM; * p ≤ 0.05, ** p ≤ 0.01, *** p ≤ 0.001 relative to 0 μM L-DOPA (A,C,D) or the corresponding 0 μM L-DOPA value at each time point (B) by paired t-test (A-D) with Holm correction for multiple comparisons (A, B, D).

Because we observed some toxicity with 200 μM L-DOPA after 24 and 48 hours of treatment (20.6 ± 1.7% cell death after 24 hours, p = 0.005, N = 4; 41.8 ± 4.9% cell death after 48 hours, p = 0.046, N = 4) (Fig. S1A), we investigated whether parkin loss can occur independently of cell death. To do this, we assessed whether subtoxic doses of L-DOPA can still decrease parkin. To maximize the chances that we would see an effect on parkin with relatively low doses of L-DOPA as well as to ensure that sufficient time would elapse to detect any cell death from L-DOPA, we used a multiple-treatment design (Fig. S1B). Differentiated PC12 cells were treated with 25-100 μM L-DOPA a total of three times, with two days between treatments. Lysates were harvested for WB two days after the final L-DOPA treatment (six days after the first exposure to L-DOPA), while replicate cultures were harvested for survival assessment three days after the final L-DOPA treatment (seven days after the first exposure to L-DOPA). In these experiments, all doses of L-DOPA were non-toxic (100.2 ± 3.9% cells remaining with 100 μM L-DOPA, p = 1.0, N = 3) (Fig. S1C), but a dose-dependent loss of parkin was again observed, reaching significance with 100 μM L-DOPA (33.9 ± 2.8% parkin loss with 100 μM L-DOPA, p = 0.03, N = 3) (Fig. 1D). These results indicate that parkin loss from L-DOPA treatment can occur independently of cell death.

Next, we determined whether L-DOPA can decrease parkin levels in cell types in addition to neuronal PC12 cells. Treatment of primary rat cortical neurons, human SH-SY5Y neuroblastoma cells, and immortalized mouse embryonic fibroblasts (MEFs) with 200 μM L-DOPA for 24 hours led to a statistically significant loss of parkin in each case (cortical neurons: 13.6 ± 5.2% parkin loss, p = 0.048, N = 6; SH-SY5Y: 32.7 ± 3.8% parkin loss, p = 0.001, N = 5; MEFs: 31.3 ± 12.0% parkin loss, p = 0.048, N = 6) (Fig. 1C; Fig. S2). Treatment of cortical neurons with a higher dose of L-DOPA, 500 μM, for 15-18 hours, led to a larger parkin loss (19.5 ± 3.3% parkin loss, p = 0.001, N = 7) (Fig. 1C). These results suggest that L-DOPA-induced parkin loss is a broadly-generalizable phenomenon and is not limited to neuronal PC12 cells.

Next, we asked whether decreased production and/or increased degradation is responsible for parkin loss after L-DOPA treatment. We did not observe a significant decrease in parkin mRNA levels after L-DOPA treatment (relative mRNA level after 27 hours normalized to 3 hours of control treatment: 0 μM L-DOPA: 1.54 ± 0.63, N = 2; 200 μM L-DOPA: 1.31 ± 0.186, N = 2; p = 1.0) (Fig. S3A), indicating that L-DOPA’s effect on parkin is not transcriptional. To assess whether L-DOPA might affect parkin turnover, we treated cells with the translation inhibitor cycloheximide (CHX) and measured parkin levels after various times with or without L-DOPA exposure (Fig. S3B). L-DOPA accelerated parkin loss in the presence of cycloheximide after an apparent lag time of several hours (CHX: 52.5 ± 5.8% parkin remaining after 48 hours, N = 3; CHX + L-DOPA: 36.2 ± 5.9% parkin remaining, N = 3; p = 0.004), such that the apparent half-life of the protein during the linear phase of decay fell from 41.9 hours to 27.2 hours without or with L-DOPA treatment, respectively. In contrast, there was no effect of L-DOPA on the halflife of ERK1 protein (Fig. S3B). These findings suggest that the loss of parkin that occurs with L-DOPA is due at least in part to its enhanced degradation.

If L-DOPA promotes parkin loss at least in part by increasing its turnover, then we would expect to see loss of overexpressed exogenous parkin. In an initial study, we did not observe consistent and specific L-DOPA-promoted loss of highly overexpressed (~30-fold) exogenous parkin driven from the EF-1α promoter (Fig. S4A), suggesting that very high levels of parkin are able to overwhelm the L-DOPA-induced parkin degradation mechanism. In contrast, when we more moderately overexpressed N-terminally-tagged parkin using the minimal human parkin promoter, leading to ~8-fold overexpression above endogenous parkin levels, we observed a L-DOPA-induced loss of exogenous parkin similar to that observed with the endogenous protein (40.7 ± 0.7% exogenous parkin loss with L-DOPA, p = 0.0003, N = 3) (Fig. S4B).

### L-DOPA-induced oxidative stress contributes to parkin loss

Given that L-DOPA has been shown to cause oxidative stress in cultured cells [56], [60], [61], we examined whether L-DOPA-induced oxidative stress plays a role in parkin loss. To do this, we treated differentiated PC12 cells with L-DOPA in the presence of the antioxidant glutathione. Glutathione (GSH) significantly attenuated parkin loss from L-DOPA, but this rescue was not complete (43.3 ± 3.9% loss with L-DOPA alone vs. 18.9 ± 6.5% loss with L-DOPA + GSH, N = 3, p = 0.03; p = 0.03 for L-DOPA + GSH vs. GSH alone, N = 3) (Fig. 2A). We did not observe enhanced protection of parkin from L-DOPA-induced loss by using higher doses of glutathione or by pre-treating cells with glutathione prior to L-DOPA exposure (Fig. S5). This result indicates that oxidative stress is at least partially responsible for L-DOPA-induced parkin loss but that a non-oxidative mechanism may account for about half of this loss.

**Figure 2.**
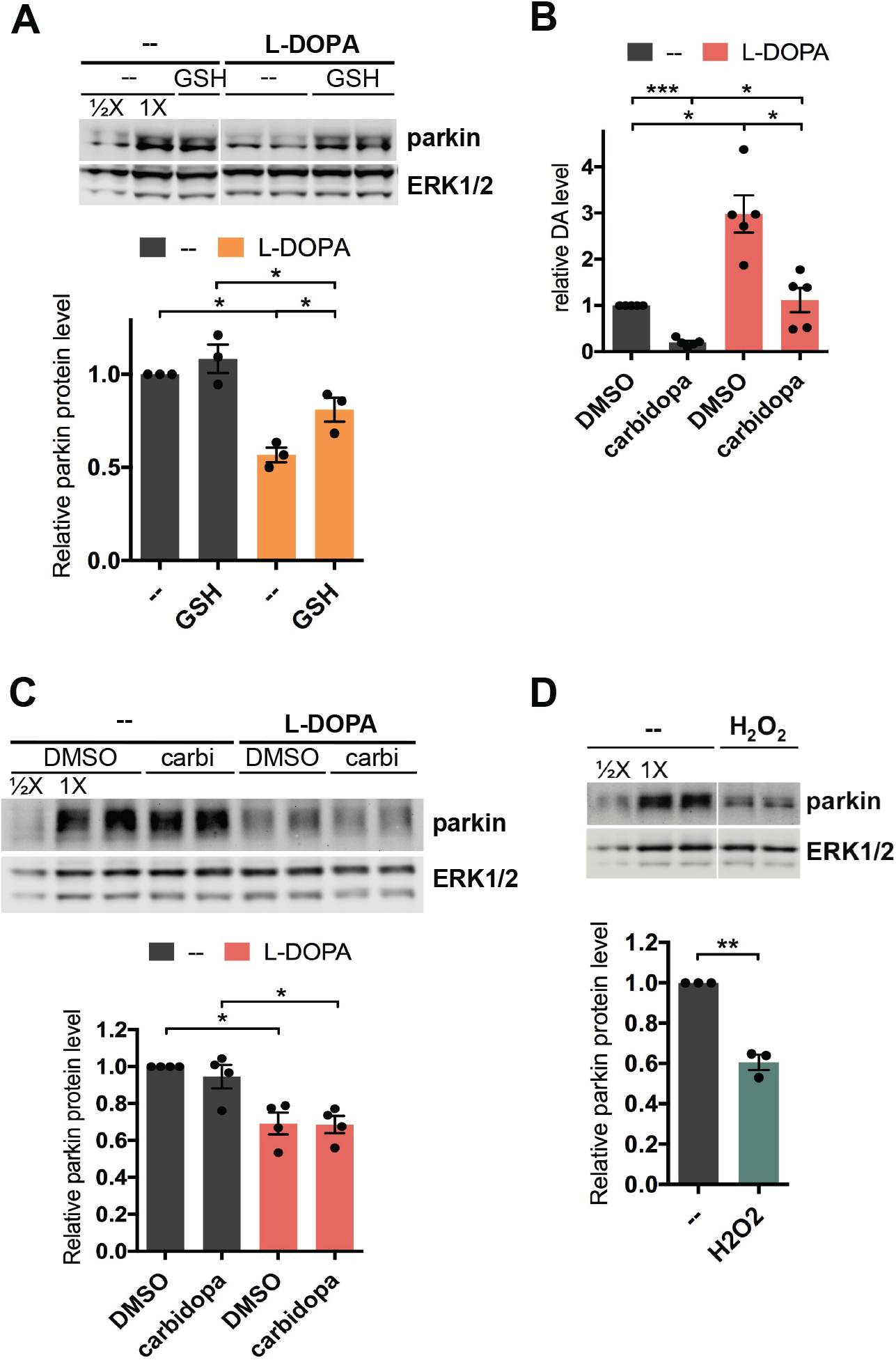
L-DOPA-induced oxidative stress contributes to parkin loss. A. Glutathione attenuates L-DOPA’s effect on parkin. Neuronally differentiated PC12 cells were cotreated with 200 μM L-DOPA and 200 μM glutathione (GSH) for 24 hours before lysates were harvested for Western immunoblotting. B,C. Treatment with carbidopa does not prevent parkin protein loss from L-DOPA treatment. Cells were pretreated with 50 μM carbidopa (carbi) for 2 hours, then co-treated with 200 μM L-DOPA. B. 16-24 hours after L-DOPA addition, cells were harvested for analysis of intracellular dopamine (DA) levels by HPLC. C. 24 hours after L-DOPA addition, lysates were harvested for Western immunoblotting. A representative Western blot and quantification of parkin levels are shown. D. Hydrogen peroxide induces parkin protein loss. Cells were treated with 200 μM hydrogen peroxide for 24 hours before lysates were harvested for Western immunoblotting. A-D. Error bars show SEM from N = 3 (A), 5 (B), 4 (C) and 3 (D) experiments; * p ≤ 0.05, ** p ≤ 0.01, *** p ≤ 0.001 by paired t-test (A-D) with Holm correction for multiple comparisons (A-C).

L-DOPA can generate reactive oxygen species (ROS) via non-enzymatic autoxidation and by conversion to dopamine, with subsequent autoxidation and oxidative metabolism of the latter [53], [62], [63]. To test whether conversion of L-DOPA to dopamine is required for its effect on parkin, we used carbidopa to block the activity of aromatic L-amino acid decarboxylase (AADC), the enzyme responsible for this conversion. As anticipated, 200 μM L-DOPA treatment alone for 16-24 hours significantly increased intracellular dopamine levels (to 3.0 ± 0.4 fold over control levels, p = 0.02, N = 5) (Fig. 2B). Co-treatment with carbidopa robustly diminished this effect (to 1.1 ± 0.3 fold over control levels, p = 0.02 vs. L-DOPA alone, N = 5) but did not impact the loss of parkin from L-DOPA treatment (30.9 ± 5.9% parkin loss with L-DOPA alone vs. 31.4 ± 4.7% loss with L-DOPA + carbidopa, p = 0.9, N = 4) (Fig. 2C). These results suggest that autoxidation of L-DOPA itself, rather than its conversion to dopamine, is the source of the oxidative stress that leads to parkin loss following L-DOPA treatment in our system. Consistent with this possibility, we observed “browning” of the cell culture medium after 24 hours of exposure to L-DOPA, even in the absence of cells (Fig. S6), which has been shown to be a result of quinone formation from L-DOPA autoxidation [61], [64]–[66].

To confirm that oxidative stress is capable of inducing parkin loss, we treated cells with 200 μM hydrogen peroxide and observed a reduction of parkin protein comparable to that found with L-DOPA (39.4 ± 3.8%, p = 0.009 vs. untreated control, N = 3) (Fig. 2D), in line with a previous report [41]. Altogether, these results indicate that oxidative stress from L-DOPA autoxidation significantly contributes to L-DOPA-induced parkin loss.

### PINK1 plays a role in L-DOPA-induced parkin loss

The kinase PINK1 has been reported to play a role in parkin turnover, including that caused by hydrogen peroxide exposure [41], [67]. We therefore examined whether PINK1 is involved in L-DOPA-dependent parkin loss. To do this, we knocked down PINK1 in differentiated PC12 cells using a previously-characterized shRNA [68]. PINK1 knockdown was very effective at the mRNA level, reducing PINK1 transcript levels to 14.1 ± 3.5% of control levels after 4 days (p = 0.0001, N = 4) (Fig. 3A).

**Figure 3.**
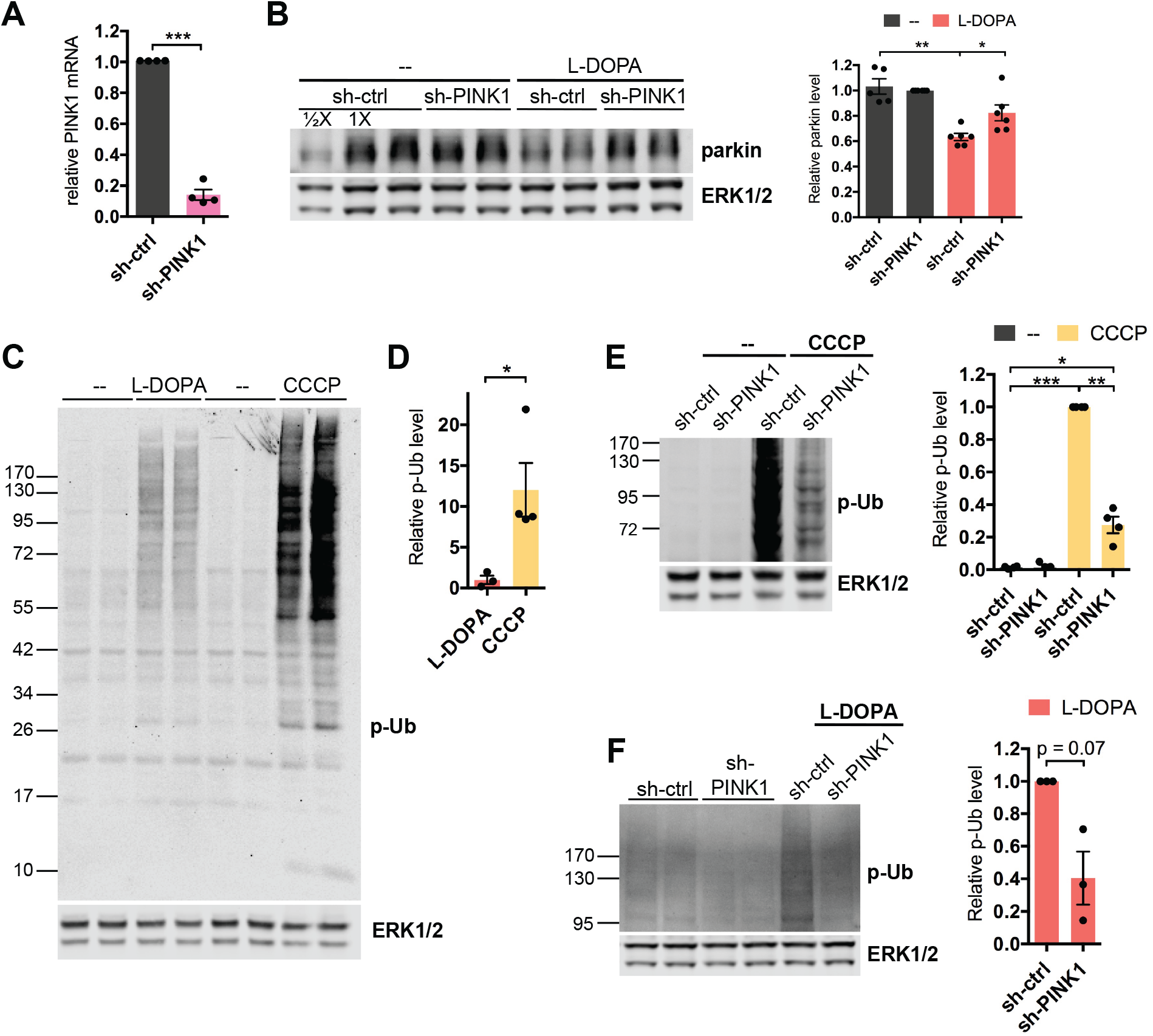
PINK1 plays a role in L-DOPA-induced parkin loss. A,B. Knockdown of PINK1 partially prevents parkin loss from L-DOPA treatment. Neuronally differentiated PC12 cells were transduced with shRNA against PINK1 for 4 days before being harvested for qPCR analysis of PINK1 mRNA levels (A) or treated with 200 μM L-DOPA (B). Cell lysates were harvested for WB after 24 hours of L-DOPA treatment (B). Representative Western immunoblots and quantification of parkin protein levels are shown. C,D. L-DOPA and CCCP induce phospho-poly-Ub formation. Differentiated PC12 cells were treated with 200 μM L-DOPA for 24 hours or 10 μM CCCP for 10 hours before being harvested for Western immunoblotting. Representative Western blot (C) and quantification (D) of phospho-poly-Ub (p-Ub) levels with L-DOPA and CCCP treatment are shown. E,F. PINK1 knockdown attenuates phospho-poly-Ub induction by L-DOPA and CCCP treatment. Cells were transduced with shRNA against PINK1 for 4 days before being treated with 10 μM CCCP (E) or 200 μM L-DOPA (F). Cell lysates were harvested for WB after 12 hours of CCCP (E) and 24 hours of L-DOPA (F) treatment. Representative blots and quantifications of p-Ub levels are shown. A-F. Error bars show SEM from N = 4 (A), 5-6 (B), 3-4 (D) 4 (E), and 3 (F) independent experiments; * p ≤ 0.05, ** p ≤ 0.01, *** p ≤ 0.001 by paired (A,B,E,F) or unpaired (D) t-test, with Holm correction for multiple comparisons (B,E).

Knockdown of PINK1 significantly attenuated L-DOPA-induced parkin loss, to about half that achieved with L-DOPA alone (36.7 ± 2.8% parkin loss with sh-ctrl vs. 17.7 ± 6.3% loss with sh-PINK1, p = 0.02 for sh-ctrl + L-DOPA vs. sh-PINK1 + L-DOPA, N = 6) (Fig. 3B). These findings indicate that PINK1 is indeed involved in L-DOPA-mediated parkin loss, but that a PINK1-independent mechanism also appears to contribute to this process.

To promote turnover of cellular proteins, PINK1 must be activated, and a role for this kinase in parkin turnover in response to L-DOPA thus suggests that the latter causes PINK1 activation. To assess this, we examined S65-phosphorylated ubiquitin (phospho-Ub) as a readout of PINK1 activity. At baseline, phospho-Ub is present at very low or undetectable levels in cells [69], [70]. Following mitochondrial depolarization, PINK1 becomes stabilized on the outer mitochondrial membrane and activated, leading it to phosphorylate ubiquitin and resulting in an accumulation of phosphorylated poly-ubiquitin chains (phospho-poly-Ub) that is detectable by Western blotting [69], [70]. As expected, we observed little phospho-Ub in untreated PC12 cells (Fig. 3C). As a positive control for PINK1 activation and phospho-Ub induction, we treated cells with the protonophore carbonyl cyanide 3-chlorophenylhydrazone (CCCP), which is widely used as a mitochondrial depolarizing agent [71]. Following CCCP treatement, we observed the appearance of high molecular weight phospho-Ub ladders, consistent with formation of phospho-poly-Ub chains (Fig. 3C). Treatment of cells with L-DOPA also led to formation of phospho-poly-Ub (Fig. 3C), although the strength of the L-DOPA-induced phospho-poly-Ub signal was about 10 times weaker than that induced by CCCP (p = 0.04, N = 3 for L-DOPA and 4 for CCCP) (Fig. 3C,D). We did not observe the formation of unconjugated mono-phospho-Ub after L-DOPA treatment (Fig. 3C). Knockdown of PINK1 strongly attenuated the phospho-poly-Ub signal from CCCP exposure (to 27.5 ± 5.1% of sh-ctrl, p = 0.003, N = 4), and caused a similar downward trend in the L-DOPA-induced phospho-poly-Ub signal (to 40.5 ± 16.3% of sh-ctrl, p = 0.07, N = 3), indicating that PINK1 indeed appears to be involved in generating this signal (Fig. 3E,F). Together, these results indicate that L-DOPA exposure leads to PINK1 activation, albeit to a lesser extent than CCCP exposure, and that this activation contributes to parkin loss.

### Association with L-DOPA-induced phospho-Ub leads to parkin loss

Having implicated PINK1 in L-DOPA-induced parkin depletion, we next sought to uncover the mechanism by which PINK1 contributes to this loss. PINK1 activates parkin by phosphorylating Ser65 on both parkin and on ubiquitin [42]–[44], [49], [72]–[77], with parkin binding the latter non-covalently. Parkin binds to both unconjugated phospho-mono-Ub and to phospho-poly-Ub chains [43], [78], [79]. We reasoned that parkin phosphorylation, parkin binding to L-DOPA-induced phospho-Ub, or both of these events may underlie PINK1-mediated parkin loss. To determine the relative importance of these events, we introduced several point mutations in the “moderately” overexpressed exogenous parkin construct described above (Fig. S4A). To test the importance of parkin phosphorylation, we generated a S65A parkin mutant. To test the importance of phospho-Ub binding to parkin, we generated parkin mutants H302A and K151E. Both of the latter mutations significantly abrogate the interaction between parkin and phospho-Ub, although K151E has been demonstrated to be more effective in this regard [47]. We delivered the parkin mutants to PC12 cells via lentiviral transduction and tested their responses to L-DOPA treatment. Although the viral titers between constructs varied slightly due to inherent variability in the viral production process, we found that all parkin mutants used in this study were expressed at similar levels (i.e. ~8-fold higher than endogenous parkin) in cells when controlling for titer (Fig. S7).

L-DOPA induced a significant loss of wild-type overexpressed parkin after 24 hours of treatment, as expected (38.0 ± 2.9% loss, p <0.0001, N = 8) (Fig. 4A,B). However, the mutants deficient in binding phospho-Ub were significantly resistant to loss compared to wild-type parkin (H302A: 22.9 ± 2.1% loss, N = 8, p = 0.006 vs. WT + L-DOPA; K151E: 20.7 ± 4.3% loss, N = 7, p = 0.003 vs. WT + L-DOPA) (Fig. 4A,B). Of note, the degree of protection from L-DOPA-induced loss afforded by the H302A and K151E mutations was ~50%, similar to the degree of protection afforded by PINK1 knockdown (Fig 3B). This suggests that the contribution of PINK1 to L-DOPA-mediated parkin loss may be fully accounted for by parkin’s interaction with PINK1-generated phospho-Ub. In agreement with this, the S65A parkin mutant was not at all protected from L-DOPA-induced loss compared to wild-type parkin (S65A: 36.8 ± 3.6% loss, N = 4, p = 0.82) (Fig. 4A,B), indicating that PINK1-mediated parkin phosphorylation is not required for parkin loss. We also tested whether a protective effect of S65A might be revealed in the context of a parkin mutant that is deficient in binding to phospho-Ub. However, we did not observe greater protection from L-DOPA with a H302A/S65A double mutant over the H302A single mutant (H302A/S65A: 16.1 ± 7.8% loss, N = 3, p = 0.48 vs. H302A alone) (Fig. 4A,B).

**Figure 4.**
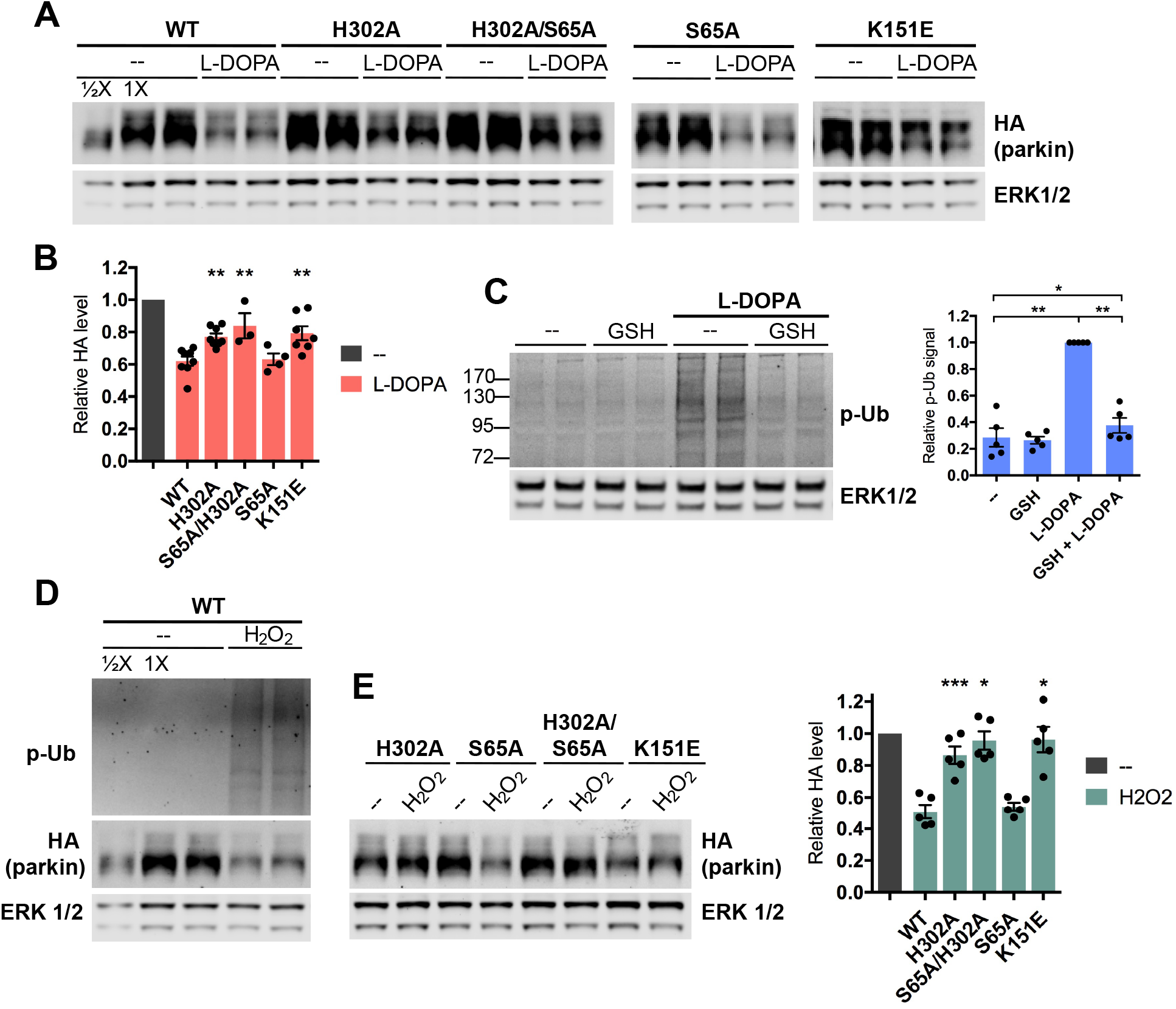
Association with oxidative stress-induced phospho-Ub leads to parkin loss. A,B. Differentiated PC12 cells were transduced with lentiviral vectors carrying the indicated parkin mutants. Three to five days after transduction, cells were treated with 200 μM L-DOPA for 24 hours before harvest for Western immunoblotting and assessment of parkin protein levels. Representative immunoblots (A) and quantifications of parkin levels (B) are shown. The level of each parkin mutant after L-DOPA treatment was normalized to the level of the same mutant after control treatment, which was set to 1 in each case. The latter is represented by the leftmost gray bar in the quantifications (B). C. Glutathione almost completely abrogates the L-DOPA-induced phospho-poly-Ub signal. Differentiated PC12 cells were co-treated with 200 μM L-DOPA and 200 μM glutathione (GSH) for 24 hours before lysates were harvested for Western immunoblotting. A representative immunoblot and quantification of phospho-poly-Ub levels are shown. D,E. Differentiated PC12 cells were transduced with lentiviral vectors as in (A) and treated with 200 μM hydrogen peroxide for 24 hours before harvest for Western immunoblotting. D. Hydrogen peroxide induces phospho-poly-Ub formation and decreases wildtype overexpressed parkin. E. Effect of hydrogen peroxide on overexpressed parkin mutants. Representative immunoblot and quantifications of parkin levels as in (B) are shown. B,C,E. Error bars show SEM from N = 3-8 (B), 5 (C), and 5 (E) independent experiments; * p ≤ 0.05, ** p ≤ 0.01, *** p ≤ 0.001 by paired t-test with Holm correction for multiple comparisons (C) or relative to WT by one-way ANOVA of stressor-treated mutants with Holm-Sidak’s multiple comparisons test (B,E).

The above findings indicate that parkin’s interaction with phospho-Ub is required for a portion of its loss following L-DOPA treatment, while parkin phosphorylation appears to be dispensable. However, one potential reservation for this interpretation is that our mutant parkin constructs have an N-terminal tag to differentiate them from endogenous parkin, and such tags have been reported to disrupt parkin’s autoinhibited conformation by opening the Ubl domain [80]. Because parkin phosphorylation is also reported to open the Ubl domain [45], [73], [81]–[84], we wished to ensure that the effect of parkin phosphorylation was not made functionally redundant by the presence of the tag. To do so, we took advantage of the finding that parkin phosphorylation is required for full parkin activation in response to CCCP [49], [77]. We compared the levels of CCCP-induced phospho-poly-Ub in cells transduced with either wild-type or S65A parkin. We reasoned that since parkin phosphorylation contributes to its activation and since such activation leads to phospho-poly-Ub formation following mitochondrial depolarization [50], we would expect to see decreased levels of phospho-poly-Ub in CCCP-treated cells expressing S65A parkin, but only if this mutant form is not already activated by the presence of the N-terminal tag. Indeed, cells transduced with S65A parkin had significantly decreased levels of phospho-poly-Ub after CCCP exposure compared with those expressing wild-type parkin (WT: 2.12 ± 0.18; S65A: 1.72 ± 0.23; p =0.04, N = 5) (Fig. S8). Similarly, CCCP-treated cells expressing the H302A/S65A double parkin mutant had significantly decreased levels of phospho-poly-Ub compared to those expressing the H302A single mutant (H302A: 1.73 ± 0.13; H302A/S65A: 1.19 ± 0.04; p = 0.007, N = 5) (Fig. S8). These results demonstrate that parkin phosphorylation plays a functional role in the CCCP model and is not made redundant by the presence of an N-terminal tag. In light of this, our observation that the S65A parkin mutant is not at all protected from L-DOPA-induced loss indicates that parkin phosphorylation status does not impact this loss. Instead, PINK1-mediated ubiquitin phosphorylation is required for PINK1-dependent parkin loss downstream of L-DOPA exposure.

### L-DOPA-induced oxidative stress- and phospho-Ub-dependent mechanisms of parkin loss are inthe same pathway

The findings that glutathione treatment, PINK1 knockdown, and mutation of parkin’s phospho-Ub binding site all rescued L-DOPA-induced parkin loss by about 50% (Figs. 2A, 3B, 4A) suggest that parkin loss from L-DOPA treatment occurs via two distinct pathways, and that oxidative stress and PINK1 activity each play a role in only one of these pathways. To determine whether the oxidative stress-dependent (glutathione-sensitive) and the phospho-Ub-dependent mechanisms of parkin loss are in the same pathway, we investigated whether glutathione can abrogate the phospho-Ub signal induced by L-DOPA. Given that the majority of this signal is present in high molecular weight phospho-poly-Ub conjugates, we examined the effect of glutathione on phospho-poly-Ub. We found that glutathione prevented L-DOPA-induced phospho-poly-Ub formation almost completely (no treatment: 28.6 ± 7.8% phospho-poly-Ub relative to L-DOPA treatment, p = 0.002, N = 5; L-DOPA +GSH: 37.6 ± 5.6% relative to L-DOPA treatment, p = 0.002, N = 5) (Fig. 4C). Additionally, hydrogen peroxide exposure induced phospho-poly-Ub formation (Fig. 4D). These results indicate that oxidative stress from L-DOPA exposure leads to PINK1 activation and that this in turn leads to the formation of phospho-Ub, its association with parkin, and parkin loss.

### Phospho-Ub generated by diverse stressors induces parkin degradation

Having placed a portion of parkin loss from L-DOPA in the same pathway as L-DOPA-induced formation of phospho-Ub, we next asked whether, and to what extent, parkin interaction with phospho-Ub plays a role in its loss from other stressors. Given that hydrogen peroxide causes parkin loss (Fig. 2D) and induces phospho-poly-Ub formation (Fig. 4D), we first queried whether parkin loss from this oxidative stressor is also mediated by its interaction with phospho-Ub. To achieve this, we utilized neuronal PC12 cells moderately over-expressing either wild-type or various mutant forms of parkin. Indeed, while wild-type parkin levels decreased by half upon peroxide treatment (50.9 ± 4.3% parkin remaining; p = 0.001, N = 5), the H302A and K151E parkin mutants were nearly completely rescued from loss induced by this agent (K151E + peroxide: 96.2 ± 8.0% parkin remaining, N = 5, p = 0.97 vs. no treatment; H302A + peroxide: 86.4 ± 5.4% parkin remaining, N = 5, p = 0.2 vs. no treatment) (Fig. 4D,E). As with the L-DOPA model, we did not observe any protection from peroxide-induced loss with the S65A mutant (S65A: 54.1 ± 2.4% parkin remaining, N = 5; p = 0.84 vs. WT) (Fig. 4E). Based on these results, it appears that parkin’s association with phospho-Ub is critical, and fully accounts for, its degradation following exposure to hydrogen peroxide.

Next, we examined whether parkin’s association with phospho-Ub is also central to its loss from the mitochondrial depolarizing agent CCCP. Mitochondrial depolarization has been shown to cause parkin loss in a PINK1-dependent manner [40]. As anticipated, CCCP led to a significant loss of wild-type overexpressed parkin after both 6 and 12 hours of treatment (33.5 ± 2.7% and 62.8 ± 1.7% parkin loss after 6 (N = 6) and 12 (N = 4) hours, respectively, p = 0.001 at both time points) (Fig. 5A,B). By contrast, both H302A and K151E parkin mutants were fully protected from CCCP-induced loss at the 6-hour time point (H302A: 95.5 ± 4.2% parkin remaining, N = 6, p = 0.66 vs. no treatment; K151E: 99.1 ± 4.3% parkin remaining, N = 5, p = 0.84 vs. no treatment) (Fig. 5A). By 12 hours, K151E remained fully protected (100.7 ± 3.7% parkin remaining, N = 3, p = 0.87 vs. no treatment), but H302A showed a loss of 26.2 ± 4.4% (N = 4, p = 0.02 vs. no treatment) (Fig. 5B). A likely explanation for the discrepancy between the behaviors of the K151E and H302A parkin mutants at 12 hours is that H302A has been shown to have greater residual phospho-Ub binding than K151E [47]. As with the L-DOPA and hydrogen peroxide models, the S65A parkin mutant was not at all protected from CCCP compared to wild-type parkin (65.9 ± 1.5% S65A (N = 5) vs. 66.5 ± 2.7% WT (N = 6) remaining at 6 hours, p = 0.9; 36.9 ± 3.2% S65A (N = 4) vs. 37.2 ± 1.7% WT (N = 4) remaining at 12 hours, p = 0.95), and we did not observe any protection of the H302A/S65A double mutant over the H302A single mutant after 12 hours of CCCP treatment (73.8 ± 4.4%H302A (N = 4) vs. 77.0 ± 0.4% H302A/S65A (N= 3) remaining, p = 0.75) (Fig. 5A,B). Altogether, these data indicate that phospho-Ub generated by diverse stressors induces parkin degradation.

**Figure 5.**
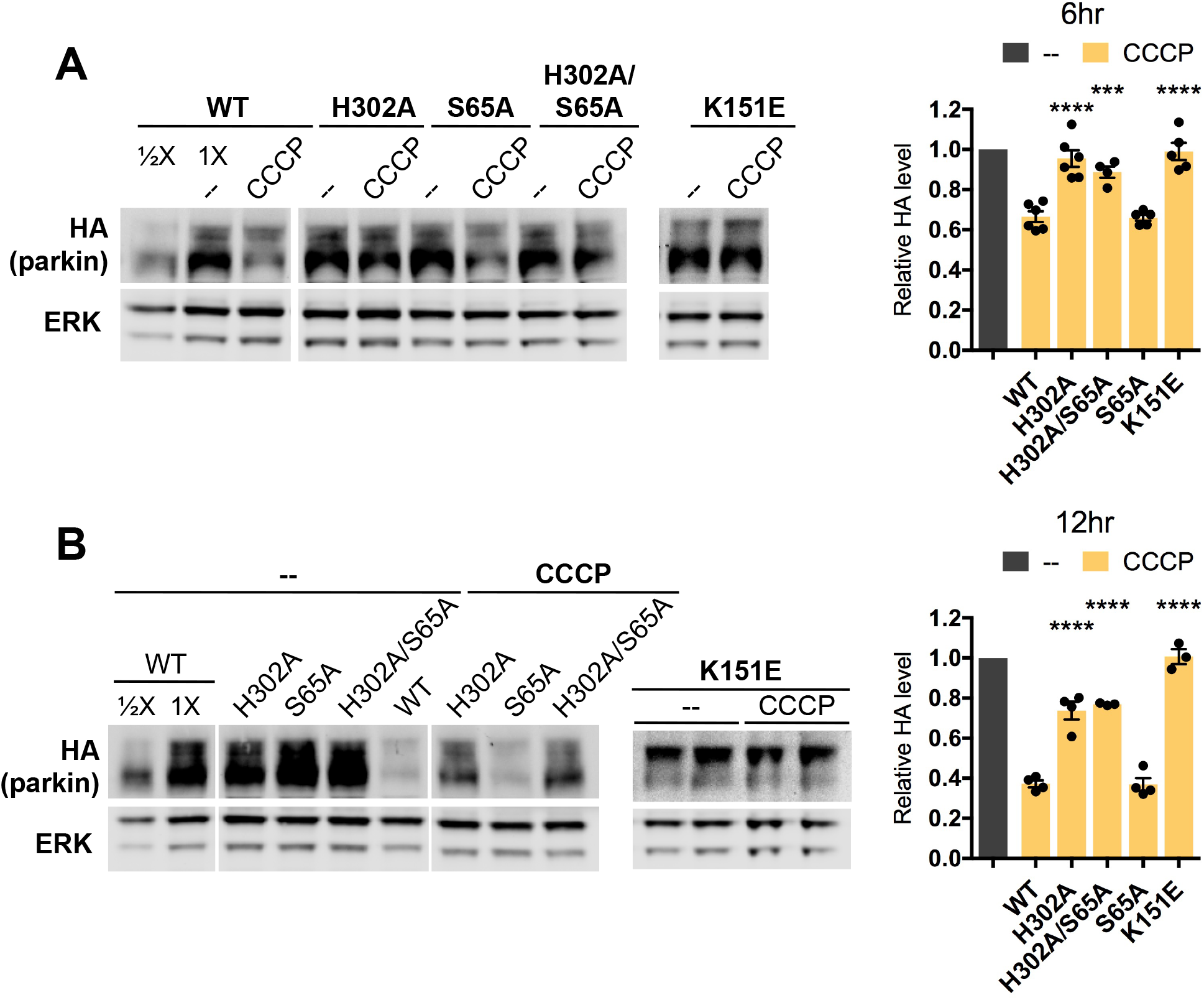
Association with phospho-Ub leads to parkin loss from CCCP treatment. A,B. Differentiated PC12 cells were transduced with lentiviral vectors carrying the indicated parkin mutants. Three to five days after infection, cells were treated with 10 μM CCCP for 6 (A) or 12 (B) hours before harvest for Western immunoblotting and assessment of parkin protein levels. Representative immunoblots and quantifications of parkin levels are shown. The level of each parkin mutant after CCCP treatment was normalized to the level of the same mutant after control treatment, which was set to 1 in each case. The latter is represented by the leftmost gray bar in the quantifications. Error bars show SEM from N = 4-6 (A) and 3-4 (B) independent experiments; *** p ≤ 0.001, **** p ≤ 0.0001 relative to WT by one-way ANOVA of CCCP-treated mutants with Holm-Sidak’s multiple comparisons test.

### Phospho-poly-Ub is present in both cytosolic and mitochondrial fractions and leads to mitochondrial parkin translocation

Given the importance of phospho-Ub for parkin loss from oxidative stress and mitochondrial depolarization, we next queried where in the cell phospho-Ub is found. As before, we examined high molecular weight phospho-poly-Ub because it is the dominant species of phospho-Ub we observe. Previous work has shown that phospho-poly-Ub is present in both the cytosol and on mitochondria following PINK1 activation by mitochondrial depolarization [69]. In line with these findings, sub-cellular fractionation revealed similar amounts of phospho-poly-Ub in the mitochondrial and cytosolic fractions after L-DOPA and CCCP treatment (L-DOPA: 1.76 ± 0.49 mitochondrial vs. cytosolic phospho-poly-Ub, p = 0.22, N = 4; CCCP: 0.81 ± 0.12 mitochondrial vs. cytosolic p-Ub, p = 0.23, N = 4) (Fig. 6A-D).

**Figure 6.**
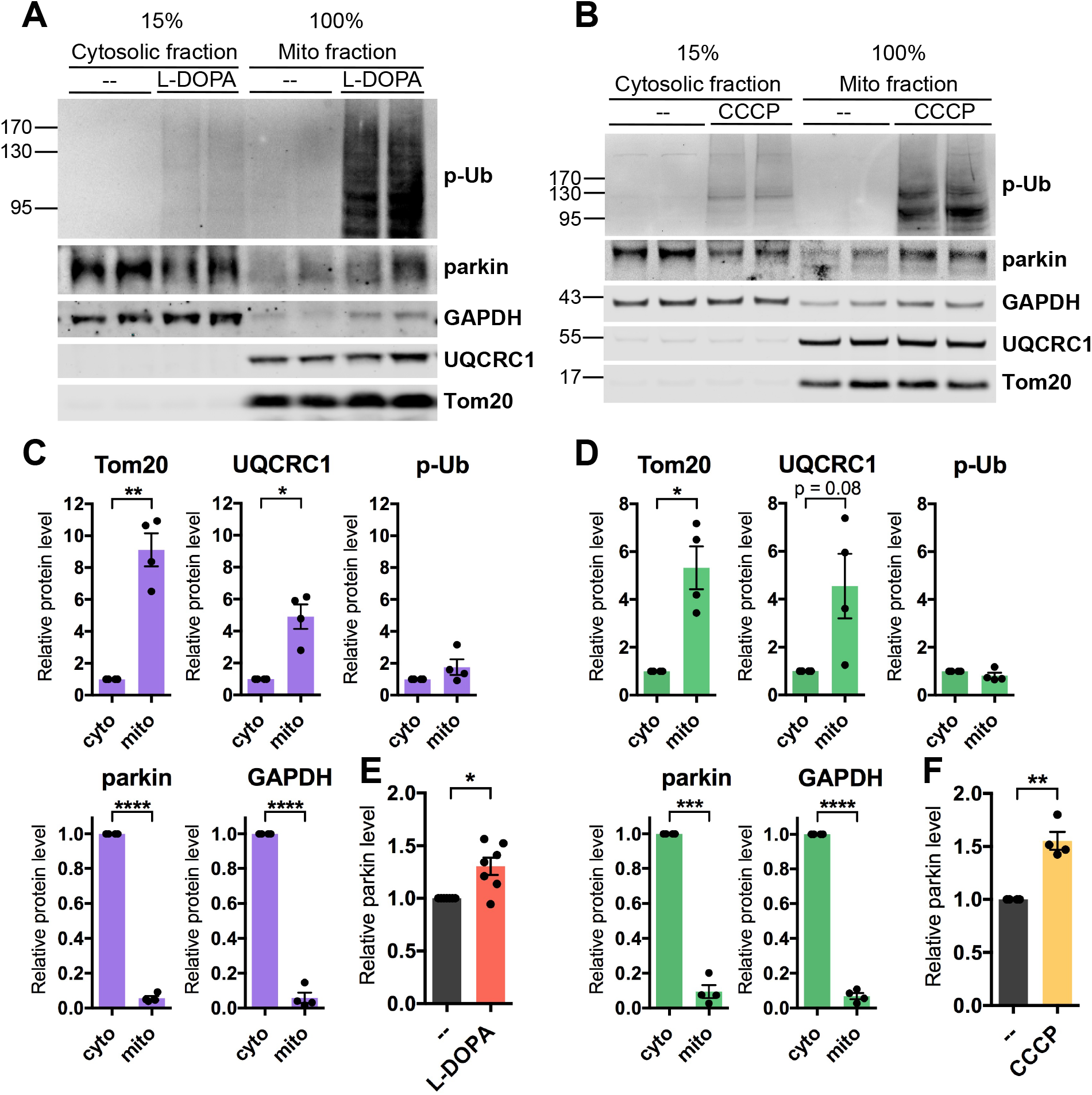
Phospho-poly-Ub is present in both cytosolic and mitochondrial fractions and leads tomitochondrial parkin translocation. A-F. PC12 cells were treated with 200 μM L-DOPA for 14-17 hours (A,C,E) or 10 μM CCCP for 6 hours (B,D,F) before being harvested for sub-cellular fractionation. 15% of the resulting cytosolic fractions and 100% of the mitochondria-enriched fractions were analyzed by Western immunoblotting for the indicated proteins. A,B. Representative Western blots showing the distribution of phospho-ubiquitin, parkin, GAPDH, UQCRC1, and Tom20 between cytosolic and mitochondrial fractions. A-D. Phospho-poly-Ubiquitin is present both in the cytosol and on mitochondria after L-DOPA (A,C) and CCCP (B,D) treatment. The phospho-poly-Ubiquitin signal in the cytosolic and mitochondrial fractions was normalized to the percentage of the fraction that was loaded to estimate the total level of phospho-poly-ubiquitin in each fraction. The same quantification was carried out for mitochondrial (Tom20, UQCRC1) and cytosolic (parkin, GAPDH) proteins from untreated cells to confirm that fractionation was successful. A,B,E,F. Parkin protein translocates to mitochondria after L-DOPA (A,E) and CCCP (B,F) treatment. Parkin levels in the mitochondrial fraction with drug or control treatment were normalized to levels of UQCRC1 and plotted in E and F. C-F. Error bars show SEM from N = 4 (C,D), 7 (E), and 4 (F) independent experiments; for F, one outlier was identified using the ROUT method with Q = 0.5% and omitted. * p ≤ 0.05, ** p ≤ 0.01, *** p ≤ 0.001, **** p ≤ 0.0001 by paired t-test.

Given the substantial presence of phospho-poly-Ub in the mitochondrial fraction after L-DOPA treatment and the affinity of parkin for phospho-Ub [47], [72], [73], [81], we predicted that we would observe mitochondrial parkin translocation following L-DOPA treatment, analogously to that observed following CCCP exposure [50], [85]. In agreement with this, we saw a modest but significant increase of parkin in the mitochondrial fraction after 14-17 hours of L-DOPA treatment (1.31 ± 0.08 with L-DOPA vs. without, p= 0.01, N = 7) (Fig. 6A,E), a time before L-DOPA-induced parkin loss is complete (Fig. 1B). As a positive control, we examined mitochondrial translocation of parkin following 6 hours of CCCP exposure - a time at which parkin loss is underway but not complete (Fig. 10B). Similarly to the effect of L-DOPA treatment, there was an increase of parkin in the mitochondrial fraction after CCCP treatment (1.55 ± 0.08 with CCCP vs. without, p = 0.007, N = 4) (Fig. 6B,F). Altogether, these results suggest that parkin associates with stress-induced phospho-poly-Ub both in the cytosol and on mitochondria before its degradation.

### Phospho-Ub-induced parkin degradation is proteasomal

We next sought to address the mechanism of parkin degradation following its association with phospho-Ub. Two previous studies reported that parkin degradation is proteasomal after mitochondrial depolarization [40], [41]. In agreement with these reports, the proteasomal inhibitor epoxomicin fully abrogated CCCP-induced parkin loss (CCCP: 63.5 ± 7.6% of untreated; CCCP + epoxomicin: 94.5 ± 7.6% of untreated, p = 0.02, N =5) (Fig. 7A). Significantly, it also abrogated L-DOPA-induced parkin loss, but by about half (L-DOPA: 61.1 ± 3.0% of untreated; L-DOPA + epoxomicin: 80.5 ± 3.3% of untreated, p = 0.001, N = 8), consistent with half of L-DOPA-induced parkin loss being phospho-Ub-dependent (Fig. 7B). Lysosomal activity and caspase cleavage have both been reported to degrade parkin [86]–[89], but inhibition of these pathways did not abrogate parkin loss from L-DOPA treatment (L-DOPA: 41.1 ± 3.5% parkin loss, L-DOPA + bafilomycin A1: 36.6 ± 1.7% parkin loss, p = 0.41, N = 3; L-DOPA: 41.0 ± 5.7% parkin loss, L-DOPA + Z-VAD-FMK: 40.8 ± 4.0% parkin loss, p = 0.98, N = 4) (Fig. S9A,B). Taken together, these data indicate that phospho-Ub-dependent parkin degradation is mediated by the proteasome in the various models tested here.

**Figure 7.**
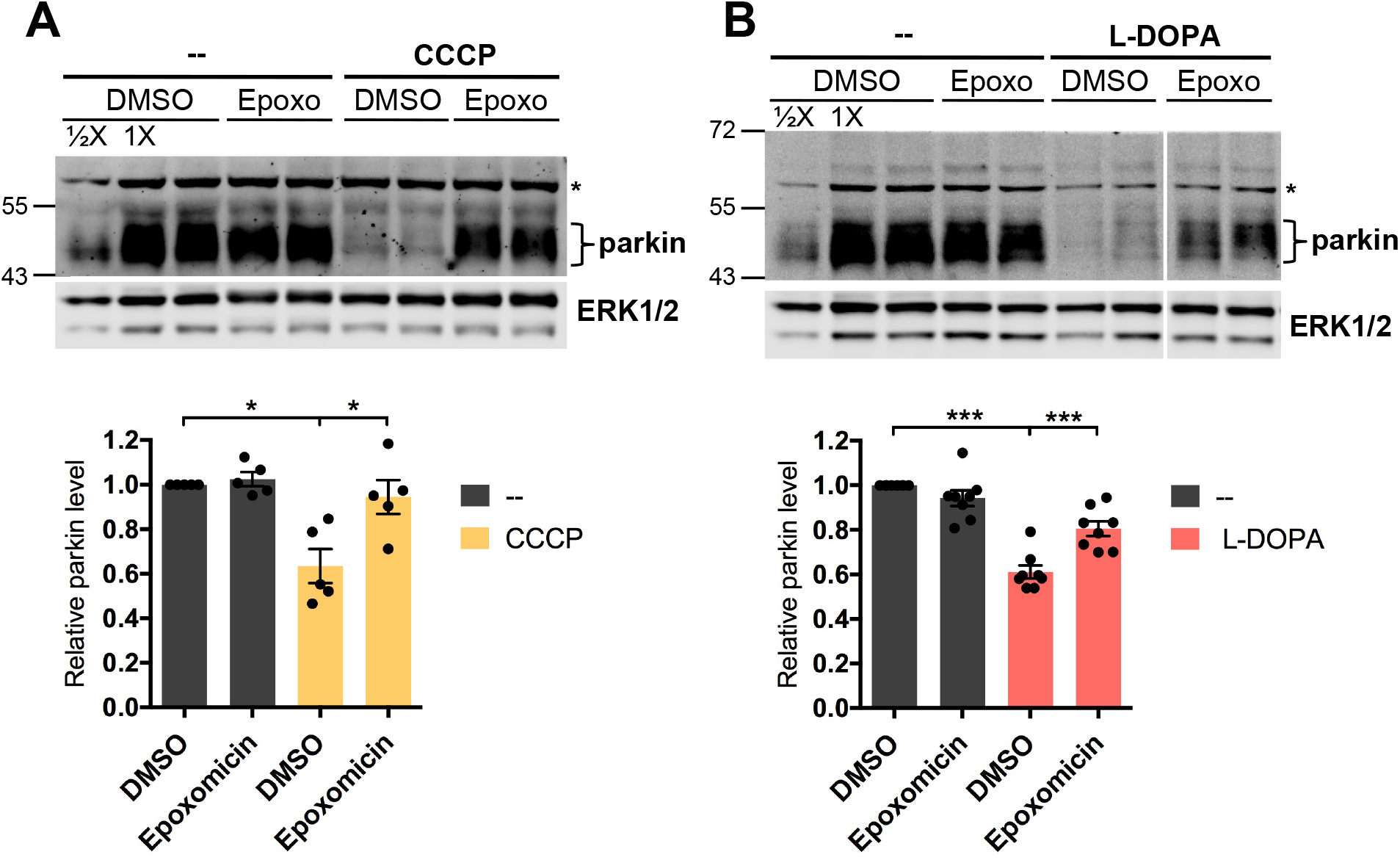
Phospho-Ub-induced parkin degradation is proteasomal. A,B. Differentiated PC12 cells were co-treated with either 10 μM CCCP (A) or 200 μM L-DOPA (B) and 100 nM epoxomicin for 12 (A) or 24 (B) hours before being harvested for Western immunoblotting. Representative Western immunoblots and quantification of parkin levels are shown. * in blots marks a known non-specific band. Error bars show SEM from N = 5 (A) and 8 (B) independent experiments; * p ≤ 0.05, *** p ≤ 0.001, by paired t-test with Holm correction for multiple comparisons.

### L-DOPA does not appear to affect parkin solubility

In addition to degradative mechanisms, various stressors have also been reported to diminish available parkin levels by disrupting its structure and decreasing its solubility [90]–[96]. To assess whether this might contribute to apparent parkin loss after L-DOPA exposure, we harvested L-DOPA-treated and non-treated cells in either our standard lysis buffer or a strong lysis buffer containing 8 M urea in addition to 2% lithium dodecyl sulfate (LDS). The strong lysis buffer did not produce significantly higher yields of parkin recovery (standard lysis buffer: 30.7 ± 4.7% parkin loss, 8 M urea buffer: 20.9 ± 4.8% parkin loss, p = 0.32, N = 5) (Fig. S10). These findings suggest that L-DOPA does not reduce apparent cellular levels of parkin by reducing its solubility.

### Phospho-Ub-dependent parkin loss does not require canonical mitochondrial parkin activity

During mitophagy initiation, PINK1 and parkin participate in a positive feedback loop wherein parkin-mediated poly-ubiquitination of mitochondrial proteins provides PINK1 with ubiquitin substrates for phosphorylation, generating phospho-poly-Ub [50]. In accordance with this process, parkin activity has been found to increase the total amount of cellular phospho-Ub following mitochondrial depolarization [69]. If this process occurred in our models, we would expect stress-induced phospho-poly-Ub formation and phospho-Ub-induced parkin loss to depend on parkin’s ligase activity.

To examine this possibility, we compared the levels of stress-induced phospho-poly-Ub in PC12 cells transduced with our “moderately overexpressed” wild-type parkin, a catalytically inactive parkin mutant (C431S), or an empty control vector. In CCCP-treated cells, expression of wild-type parkin increased the phospho-poly-Ub signal by about 3-fold over that of cells expressing empty vector, as anticipated (WT: 2.85 ± 0.30 relative to empty vector, p = 0.01, N = 5) (Fig. 8A,B). This increased phospho-poly-Ub signal can be attributed to parkin activity because catalytically inactive parkin almost completely abrogated the increased phospho-poly-Ub signal (C431S: 1.31 ± 0.07, p = 0.01 for WT vs. C431S, N = 5) (Fig. 8A,B). Unexpectedly, and by contrast, the L-DOPA-induced phospho-poly-Ub signal was not detectably increased in cells expressing wild-type parkin compared to empty vector or C431S parkin (WT: 0.83 ± 0.02, p = 0.04 vs. empty vector; C431S: 1.25 ± 0.12, p = 0.19 for WT vs. C431S, N = 3) (Fig. 8D,E). A similar result was observed in the hydrogen peroxide model (WT: 1.19 ± 0.17, p = 0.66 vs. empty vector; C431S: 0.92 ± 0.13, p = 0.51 for WT vs. C431S, N = 5) (Fig. 8G,H). These results suggest that parkin is not the ligase that builds the majority of the poly-ubiquitin chains that become phosphorylated by PINK1 in the L-DOPA and hydrogen peroxide models, in contrast to the positive feedback loop observed with CCCP.

**Figure 8.**
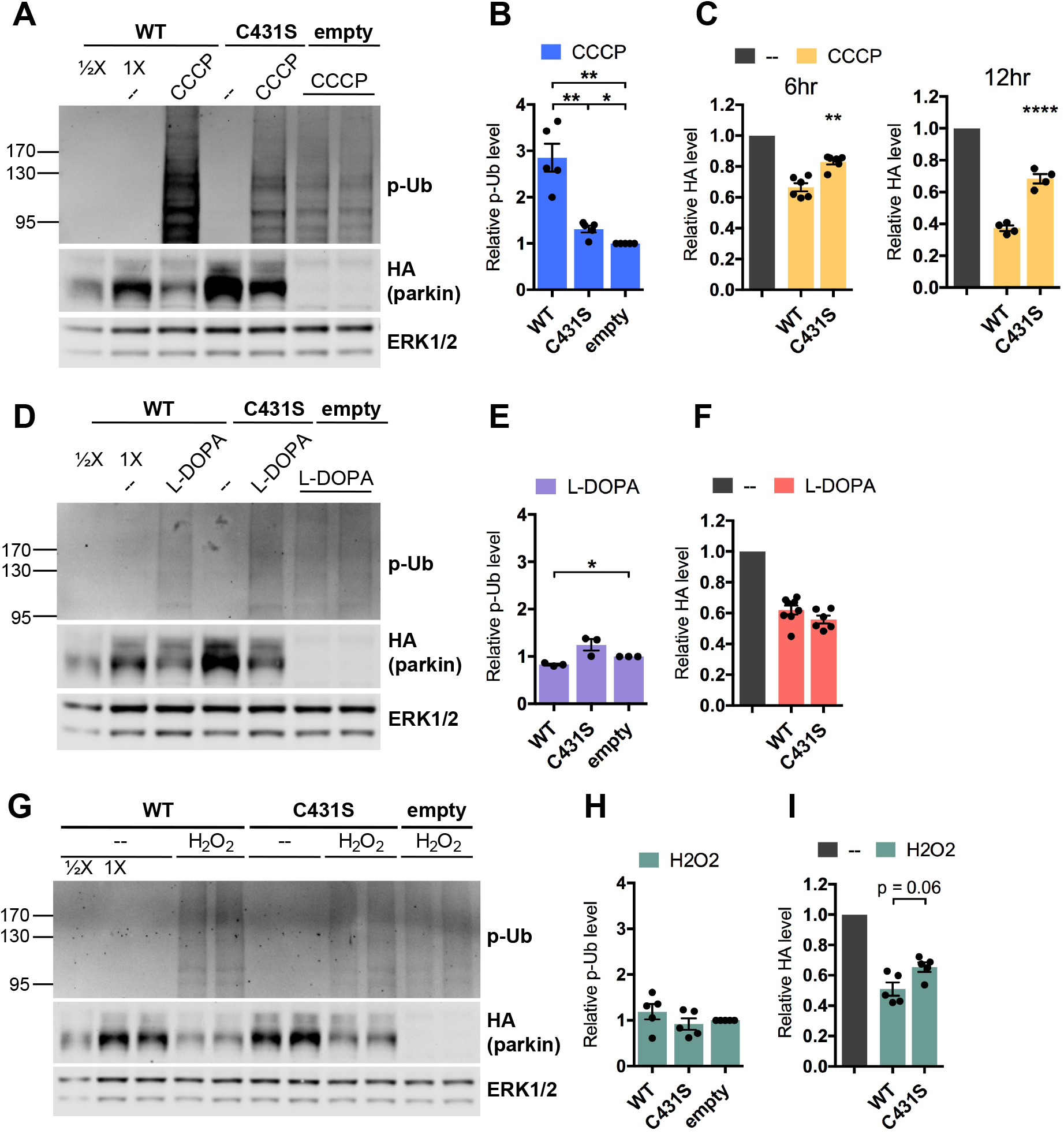
Parkin-mediated phospho-poly-Ub formation appears unnecessary for phospho-Ub-dependent parkin loss. A-I. Differentiated PC12 cells were transduced with lentiviral vectors carrying wild-type or C431S parkin. Three to five days after transduction, cells were treated with 10 μM CCCP for 6 hours (A-C) or 12 hours (C), 200 μM L-DOPA for 24 hours (D-F), or 200 μM hydrogen peroxide for 24 hours (G-I) before being harvested for Western immunoblotting. Representative Western immunoblots and quantifications of phospho-poly-Ub (B,E,H) and parkin (C,F,I) levels are shown. Error bars show SEM from N = 5 (B), 6 (C, left), 4 (C, right), 3 (E), 6-8(F), 5 (H,I) independent experiments; * p ≤ 0.05, ** p ≤ 0.01, **** p ≤ 0.0001 by paired t-test with Holm correction for multiple comparisons (B,E,H) or relative to WT by one-way ANOVA of drug-treated mutants with Holm-Sidak’s multiple comparisons test (C,F,I). The leftmost half of (G) also appears in Fig. 4C.

The lack of evidence for parkin-dependent phospho-poly-Ub formation following L-DOPA and hydrogen peroxide treatment suggested that mitochondrial parkin activity is not necessary for its phospho-poly-Ub-dependent loss. Indeed, L-DOPA exposure decreased catalytically inactive C431S parkin levels to the same extent as it did with wild-type parkin (WT: 38.0 ± 2.9% loss, N = 8; C431S: 44.2 ± 2.5% loss, N = 6; p = 0.48) (Fig. 8D,F). By contrast, C431S parkin was protected from CCCP-induced loss by ~50% after both 6 and 12 hours (6 hr: WT: 33.5 ± 2.7% loss, C431S: 16.9 ± 1.8% loss, p = 0.002, N = 6; 12 hr: WT: 62.8 ± 1.7% loss, C431S: 31.7 ± 3.0% loss, p <0.0001, N = 4) (Fig. 8A,C), consistent with the diminished phospho-poly-Ub signal observed in C431S-expressing cells relative to cells expressing exogenous wild-type parkin. In the case of hydrogen peroxide, C431S was modestly protected from peroxide-induced loss (WT: 49.1 ± 4.3% loss, C431S: 34.6 ± 3.2% loss, p = 0.06, N = 5), an effect that was less robust than that seen with CCCP. These results indicate that the extent to which parkin-mediated phospho-poly-Ub formation plays a role in parkin loss depends on the nature of the initiating stress.

The lack of evidence for phospho-poly-Ub formation by exogenous parkin following L-DOPA treatment suggests that endogenous parkin similarly does not contribute to the formation of these chains in the L-DOPA model. To examine this possibility for mitochondrial phospho-poly-Ub, we assessed ubiquitination of the mitochondrial GTPase mitofusin-2 (Mfn2) following L-DOPA treatment. A recent study showed that Mfn2 is a preferred mitochondrial parkin substrate during mitophagy and provided evidence that Mfn2 poly-ubiquitination by parkin precedes and may be required for parkin to efficiently ubiquitinate other mitochondrial substrates [97]. This primacy of Mfn2 ubiquitination, coupled with the finding that poly-ubiquitin chains on mitochondrial proteins are phosphorylated to generate mitochondrial phospho-poly-Ub, makes Mfn2 ubiquitination a useful proxy for the contribution of endogenous parkin to mitochondrial phospho-poly-Ub formation.

We first assessed Mfn2 ubiquitination following CCCP treatment, which revealed rapid poly-ubiquitination and ultimate loss of the former (32.4 ± 4.5% loss after 25 hours relative to time zero, p = 0.004 vs. no treatment, N = 3) (Fig. 9A,B). By contrast, we did not observe any ubiquitination or loss of Mfn2 after up to 48 hours of L-DOPA treatment despite robust (~50%) parkin loss (115.9 ± 4.5% Mfn2 remaining after 48 hours relative to time zero, p = 0.77 vs. no treatment, N = 3) (Fig. 9C,D; Fig. 10A). This result is in line with our findings using overexpressed parkin and indicates that phospho-Ub-dependent degradation of endogenous parkin does not require parkin’s canonical mitochondrial poly-ubiquitination activity.

**Figure 9.**
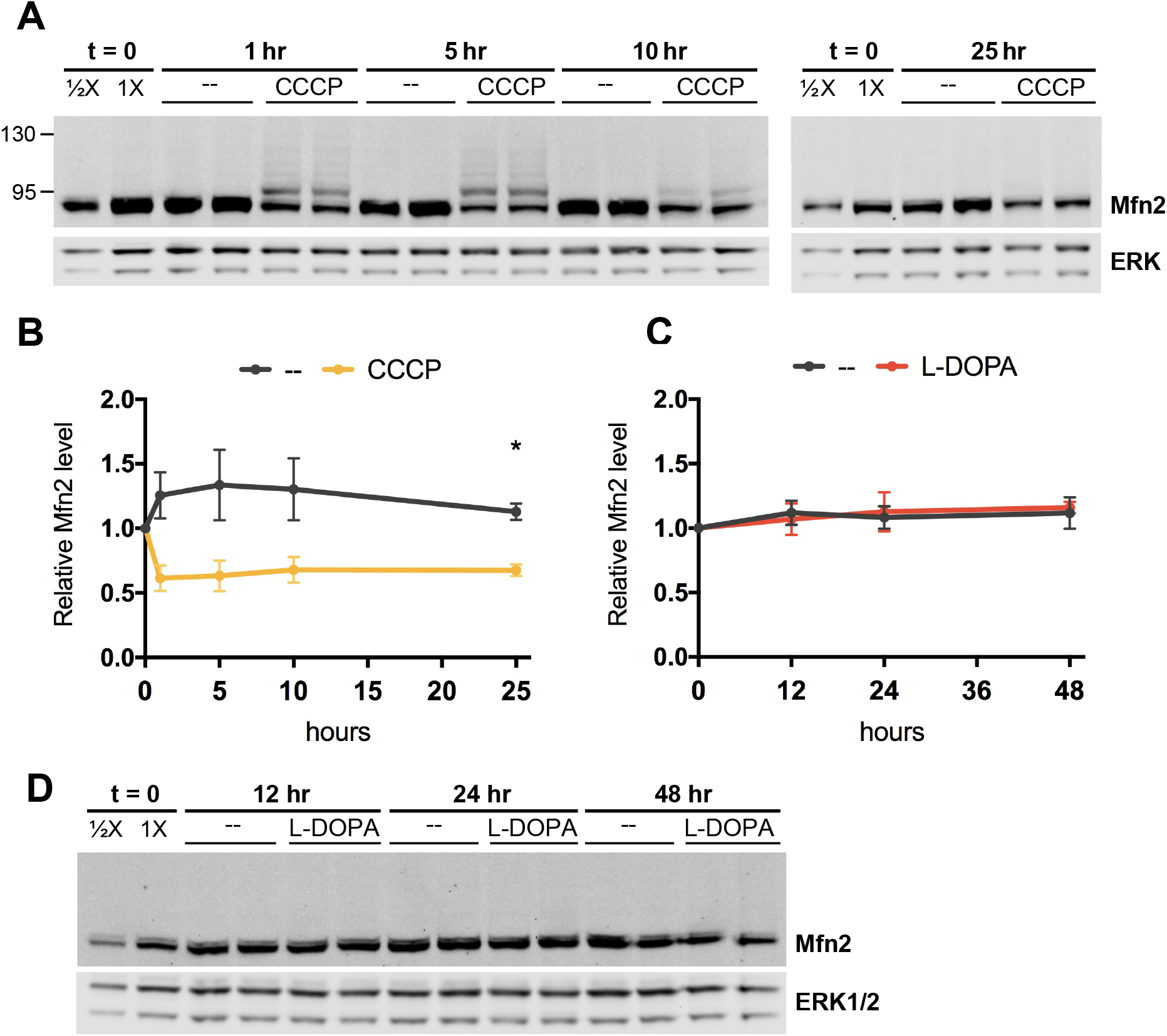
Parkin does not poly-ubiquitinate Mfn2 following L-DOPA treatment. A-D. Differentiated PC12 cells were treated with 10 μM CCCP (A,B) or 200 μM L-DOPA (C,D) for the indicated time periods before harvest for Western immunoblotting. Representative immunoblots (A,D) and quantification of unmodified Mfn2 levels from N = 3 independent experiments (B,C) are shown. Error bars show SEM; * p ≤ 0.05 by unpaired t-test with Holm-Sidak correction for multiple comparisons. Significance was queried between untreated and stressor-treated groups at each time point.

**Figure 10.**
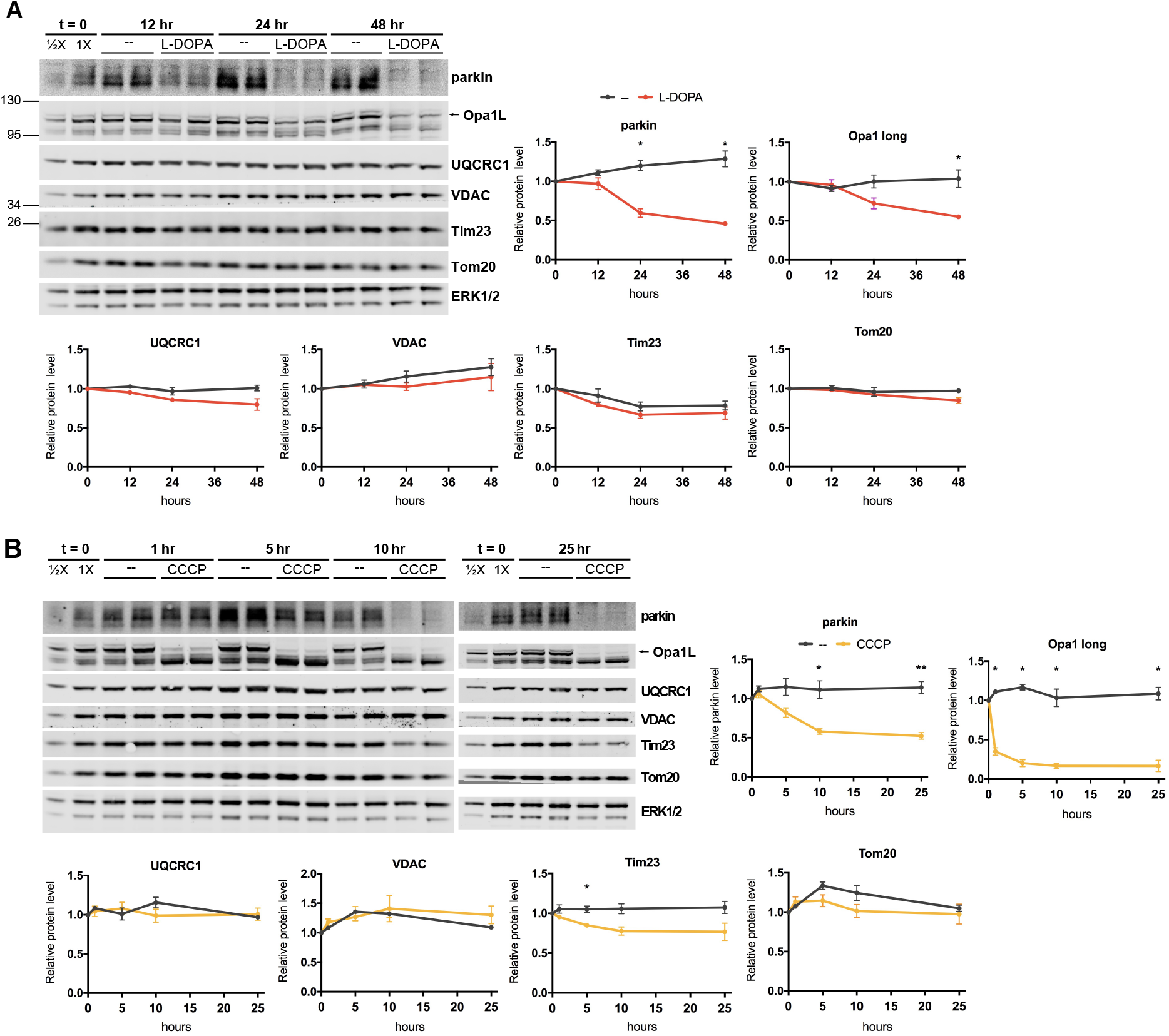
Mitophagy is not required for phospho-Ub-dependent parkin degradation. A,B. L-DOPA and CCCP do not induce changes in mitochondrial protein levels consistent with robust mitophagy. Differentiated PC12 cells were treated with 200 μM L-DOPA (A) or 10 μM CCCP (B) for the indicated times before harvest for WB. Representative blots and quantifications of the indicated proteins from N = 3 independent experiments are shown. A. L-DOPA does not significantly decrease levels of UQCRC1, VDAC, Tim23, or Tom20, while it does induce loss of parkin and Opa1-long. B. CCCP does not significantly decrease levels of UQCRC1, VDAC, or Tom20, while it does induce loss of parkin, Tim23, and Opa1-long. Error bars show SEM; * p ≤ 0.05 by unpaired t-test, with Holm-Sidak correction for multiple comparisons. Significance was queried between untreated and drug-treated groups at each time point.

### Phospho-Ub-dependent parkin loss does not require parkin autoubiquitination

Of note, given that parkin has been shown to only autoubiquitinate in cis [98], the finding that C431S parkin was not protected from L-DOPA-induced depletion indicates that this loss is not due to autoubiquitination. Parkin autoubiquitination has been reported following mitochondrial depolarization, reflected by the formation of higher-molecular weight parkin species as detected by WB [36], [40], [99]–[101]. It was previously suggested that such autoubiquitination is what targets parkin for proteasomal degradation following mitochondrial depolarization [40]. In the studies where parkin autobiquitination was observed, the ubiquitinated parkin signal was generally strongest for the band corresponding to mono-ubiquitinated parkin, but multiply-ubiquitinated parkin species were also detected, though with diminished intensity. To extend our observations with C431S parkin, we treated cells with either L-DOPA or CCCP in the presence or absence of epoxomicin and carried out Western blotting to detect higher-molecular weight parkin forms (Fig 7A,B). In no case did we detect higher-molecular weight parkin bands, consistent with the idea that autoubiquitination is not required for parkin loss.

### Phospho-Ub-dependent parkin loss does not require mitophagy

The lack of ubiquitination and loss of Mfn2 upon L-DOPA treatment suggested that L-DOPA does not induce parkin-mediated mitophagy. To determine whether this is the case, we assessed the levels of other mitochondrial proteins after L-DOPA treatment. Mitophagy leads to a gradual decrease in mitochondrial proteins in all compartments, as detected by WB [36], [102], [103]. However, we did not observe ubiquitination or a significant loss of any of the mitochondrial proteins we evaluated, including VDAC (OMM), Tom20 (OMM), Tim23 (IMM), and UQCRC1 (IMM/matrix), through 48 hours of L-DOPA treatment (Fig. 10A). To confirm the efficacy of L-DOPA treatment in these experiments, we assessed parkin levels and observed the anticipated decrease (54.1 ± 2.2% loss after 48 hours relative to time zero, p = 0.001 vs. no treatment, N = 3) (Fig. 10A). In addition, we assessed levels of the long isoform of OPA1, a mitochondrial GTPase that is cleaved in response to mitochondrial stress [104]. L-DOPA treatment induced loss of this isoform (45.0 ± 0.6% decrease after 48 hours relative to time zero, p = 0.01 vs. no treatment, N = 3) (Fig. 10A), consistent with a promotion of mitochondrial stress. Taken together, our data indicate the presence of mitochondrial stress, but a lack of robust mitophagy in the L-DOPA model.

Given that parkin was clearly active in PC12 cells after CCCP treatment (Fig. 8A,B; Fig. 9A,B), we also sought to determine whether this activity led to mitophagy in CCCP-treated PC12 cells. Examining the same proteins as we did for L-DOPA treatment, we did not observe ubiquitination or loss consistent with mitophagy in cells exposed to CCCP (Fig. 10B). Levels of Tim23 did decrease with CCCP (15.1 ± 2.6% decrease after 5 hours relative to time zero, p = 0.01 vs. no treatment, N = 3), but those of VDAC, Tom20, and UQCRC1 remained unchanged (Fig. 10B). As with L-DOPA treatment, parkin loss was still robust in these experiments (47.6 ± 4.3% loss after 25 hours, p = 0.002 vs. no treatment, N = 3) (Fig. 10B), and levels of the long isoform of OPA1 dropped precipitously in response to the mitochondrial membrane potential loss (83.5 ± 7.1% decrease after 25 hours relative to time zero, p = 0.001 vs. no treatment, N = 3) (Fig. 10B). These results indicate that even when parkin is clearly active and Mfn2 has been poly-ubiquitinated, mitophagy may not necessarily take place, consistent with a previous report [40]. Altogether, our data indicate that phospho-Ub-mediated parkin degradation does not require canonical mitochondrial parkin activity or mitophagy.

## Discussion

In this study, we investigated the mechanism by which stressors, L-DOPA in particular, decrease cellular levels of parkin protein. We found that L-DOPA causes parkin loss via two distinct pathways: an oxidative stress-dependent pathway driven by L-DOPA autoxidation and an oxidative stress-independent pathway, each of which is responsible for about half of parkin loss (Fig. 11). We conclude that oxidative stress plays a role in only one of these pathways because parkin mutants deficient in binding phospho-Ub were only partly protected from L-DOPA-induced loss, whereas they were fully protected from loss induced by the oxidative stressor hydrogen peroxide. Furthermore, exposure to the antioxidant glutathione, which is critical for neuronal defense against ROS [105], only attenuated L-DOPA-induced parkin loss by half. Although glutathione lacks membrane permeability [106], [107] and does not react efficiently with all kinds of ROS [108], glutathione treatment has been previously shown to be effective at completely preventing the autoxidation of both L-DOPA and the dopamine analog 6-hydroxydopamine (6-OHDA) to their respective quinones [109], [110]. Additionally, glutathione treatment has been found to completely prevent cell death induced by L-DOPA [109], 6-OHDA [110], dopamine [111]–[113], hydrogen peroxide [110], and the oxidant nitric oxide [114]. In support of glutathione’s capacity to prevent L-DOPA-induced oxidative stress in our hands, we observed an almost complete abrogation of L-DOPA-induced phospho-poly-Ub formation upon glutathione treatment. Nevertheless, it’s possible that not all aspects of L-DOPA-induced oxidative stress were fully neutralized by glutathione in our model, which would have led us to underestimate the contribution of oxidative stress to parkin loss.

**Figure 11.**
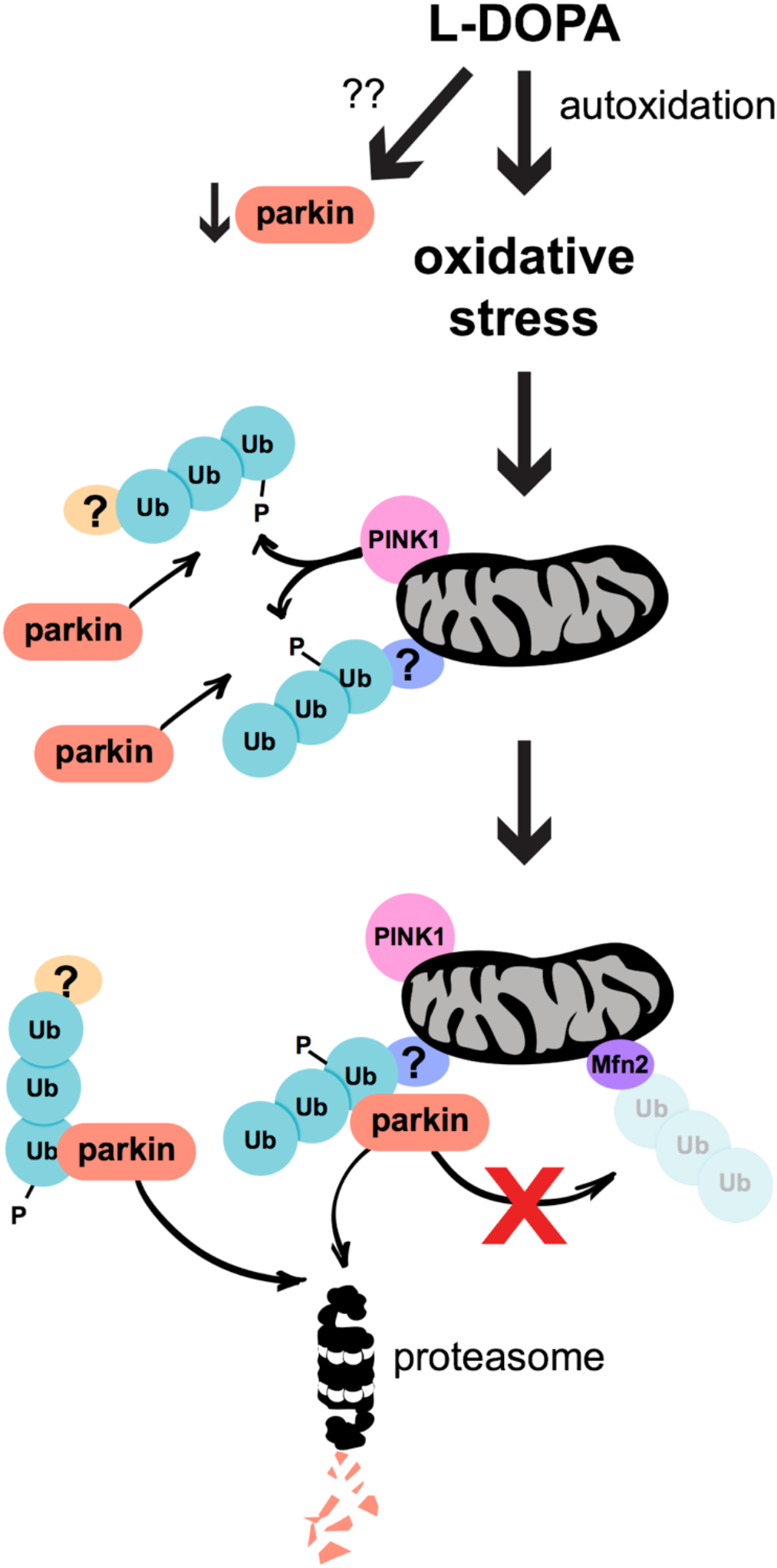
Model of L-DOPA-induced parkin loss. L-DOPA decreases parkin via two pathways: oxidative stress-dependent and -independent. The mechanism of the latter pathway remains unclear. The former pathway results from L-DOPA autoxidation, which induces PINK1 stabilization on the mitochondrial outer membrane. PINK1 phosphorylates poly-ubiquitin chains in both the cytosol and on mitochondria, though it’s unclear to which substrate protein(s) these poly-ubiquitin chains are conjugated (indicated by single question marks in the model). Parkin associates with these phospho-poly-Ub chains, but does not itself build ubiquitin chains on mitochondrial proteins. Following its association with phospho-poly-Ub, parkin is degraded via the proteasome independently of autoubiquitination and mitophagy.

Because oxidative stress has been well-documented in PD [32]–[34], we focused on the oxidative stress-dependent pathway of parkin loss. We found that, in this pathway, oxidative stress leads to PINK1-mediated ubiquitin phosphorylation, mostly of high molecular weight poly-ub conjugates. Furthermore, we found that parkin’s association with phospho-Ub induced by various stressors leads to its proteasomal degradation, but not through autoubiquitination. Surprisingly, phospho-Ub-induced parkin degradation did not appear to require parkin’s mitochondrial activity.

Our experiments with parkin point mutants reveal that parkin binding to stress-induced phospho-Ub is critical for its PINK1-dependent loss, while parkin phosphorylation, per se, is dispensable. The finding that S65A parkin was not protected from loss induced by any of the stressors we used was unexpected because phosphorylation of S65 increases the binding affinity of parkin for phospho-Ub [46], [47], [79], [81]. Given the importance of phospho-Ub binding for parkin loss, we anticipated that S65A parkin would exhibit greater protection from stress-induced degradation due to a lower affinity for phospho-Ub. On the contrary, both wild-type and S65A parkin were equally depleted under the conditions of our studies, though we cannot rule out the possibility that wild-type parkin is lost more rapidly in response to stress than the S65A mutant.

An important question that our work raises is how parkin binding to phospho-Ub following cellular stress leads to its loss. One possibility is that the conformational change in parkin that occurs upon binding phospho-Ub [45]–[47], [81], [115] makes it vulnerable to degradation. If this is the case, the conformational change must be specific to phospho-Ub binding and not to that induced by parkin phosphorylation, because we show that only phospho-Ub binding is important for parkin loss. Both events have been proposed to release parkin’s Ubl domain from its interaction with the RING1 domain [45]–[47], [81], [116], and both have also been shown to facilitate the access of E2 ubiquitin-conjugating enzymes to parkin’s E2 binding site [46], [74]. Accordingly, neither of these effects is likely to be the crucial conformational change involved in parkin degradation. Instead, a conformational change that may promote parkin loss is movement of the IBR domain away from the Ubl domain. This movement has, thus far, only been attributed to phospho-Ub binding [47], [115], [117], [118], raising it as a plausible candidate trigger for parkin loss. Alternatively, instead of inducing a critical conformational change, parkin’s association with phospho-Ub may cause parkin loss by bringing it into proximity with proteins that promote its degradation.

Our finding that the proteasome mediates parkin degradation downstream of its association with phospho-Ub is in line with prior work that reported proteasomal parkin degradation following mitochondrial depolarization [41], [119] or PINK1 overexpression [67]. Additionally, our results support the findings of others that parkin can be degraded proteasomally [15], [92], [120]–[122]. The latter and additional studies [98], [123]–[127] suggested that proteasomal parkin degradation is mediated by autoubiquitination, but this possibility was not directly tested. Our observation that catalytically inactive parkin was not protected from L-DOPA-induced loss indicates that parkin’s proteasomal degradation downstream of stress-induced phospho-Ub binding is not due to autoubiquitination, as this only occurs in cis [98]. This finding is contrary to a model proposed to explain depolarization-induced parkin loss [40], but is consistent with the finding that parkin autoubiquitination appears to protect it from degradation [36].

An additional finding of interest is that phospho-Ub-dependent parkin degradation does not appear to require parkin’s mitochondrial poly-ubiquitination activity, since Mfn2 poly-ubiquitination and loss did not occur in response to L-DOPA, despite robust parkin loss, and since parkin overexpression did not increase phospho-poly-Ub formation. This was unexpected because PINK1 and parkin participate in a well-established positive feedback loop that relies on the activity of both enzymes to generate phospho-poly-Ub during mitophagy initiation [50]. These results therefore indicate that parkin may associate with phosphorylated poly-Ub chains generated by another/other ligases.

We also observed that treatments with L-DOPA or CCCP that promoted parkin loss did not cause a loss of mitochondrial proteins consistent with robust induction of mitophagy. These results therefore indicate that mitophagy is not required for phospho-Ub-dependent parkin degradation. This finding is inconsistent with suggestions that parkin degradation downstream of mitochondrial depolarization is linked to mitochondrial elimination [36]–[39]. Instead, our results indicate that parkin’s association with phospho-Ub is the key determinant of its depolarization- and oxidative-stress-induced loss and that this loss can occur independently of mitophagy, in line with a previous study [40].

An important implication of our finding that phospho-Ub is involved in parkin loss is that a parkin activator (phospho-Ub) appears to be intimately linked with parkin degradation. We observed this phenomenon across different models of cellular stress, suggesting that it is broadly generalizable. Therefore, it is possible that degradation is a cellular mechanism by which activated parkin is “turned off”.

A recent study found that phospho-Ub levels are elevated in the substantia nigra of patients with Lewy body disease, a category that includes PD [128]. Given our findings, this observation suggests that parkin levels may be lowered in these patients due to enhanced phospho-Ub-induced turnover. This decrease could in turn contribute to PD pathology. As such, finding ways to uncouple parkin’s association with phospho-Ub from its degradation may be an attractive therapeutic avenue for PD treatment.

## Materials & Methods

### Antibodies

| Target | Dilution | Company | Catalog Number |
| --- | --- | --- | --- |
| Cleaved caspase 3 | 1:1000 | Cell Signaling Technology | 9661 |
| ERK1/2 | 1:5000 | Santa Cruz Biotechnology | SC-93 |
| GAPDH | 1:1000 | Novus Biologicals | NB 300-221 |
| GFP | 1:1000 | Thermo Fisher Scientific | A-11122 |
| HA | 1:1000 | Cell Signaling Technology | 3724 |
| LC3B | 1:1000 | Abcam | ab48394 |
| Mfn2 | 1:1000 | Cell Signaling Technology | 9482 |
| Opa1 | 1:1000 | Cell Signaling Technology | 80471 |
| Parkin | 1:500 | Cell Signaling Technology | 4211 |
| Parkin | 1:500 | Cell Signaling Technology | 2132 |
| phospho-Ub | 1:1000 | Cell Signaling Technology | 37642 |
| Tim23 | 1:1000 | BD Biosciences | 611222 |
| Tom20 | 1:1000 | Cell Signaling Technology | 42406 |
| UQCRC1 | 1:2000 | Thermo Fisher Scientific | 459140 |
| VDAC | 1:1000 | Cell Signaling Technology | 4866 |
| Mouse | 1:5000 | LI-COR | 925-68070 |
| Rabbit | 1:5000 | LI-COR | 925-32211 |

### Cell culture

### PC12 cells

Cells were grown on dishes coated with rat tail collagen (Roche; Millipore Sigma). Collagen was dissolved in 5 mL 0.2% (v/v) acetic acid and diluted 1:20 in sterile, deionized water prior to coating plates. 139 μL/cm^2^ or 17.6 μL/cm^2^ of diluted collagen was added to multiwell plates or 10-cm dishes, respectively, and plates were left uncovered to dry overnight in a laminar flow hood. Cells were plated the following day.

Undifferentiated PC12 cells were maintained in RPMI 1640 (Corning) supplemented with 10% horse serum (Millipore Sigma), 5% fetal bovine serum (Gemini BioProducts), and 1% penicillin-streptomycin. When cells were ≥70% confluent, they were split or used for differentiation.

To differentiate PC12 cells, undifferentiated cells from one 10-cm dish were first resuspended in 4mL RPMI with 1% horse serum and 1% penicillin-streptomycin. Then, cells were passed through a 5-mL repeat pipettor tip to disperse them, counted using a hemacytometer, and plated at a density of ~3×10^5^ cells/mL (1mL/well in a 12-well plate) in RPMI with 1% horse serum, 1% penicillin-streptomycin, and 50 ng/mL human recombinant NGF (kind gift of Genentech, Gemini Bio-Products #300-174P). The remaining undifferentiated PC12 cells were plated on a freshly-coated 10 cm dish. Cells were used in experiments after 5-10 days of differentiation. In experiments with lentiviral transduction, cells were transduced after 3-5 days of differentiation; for shRNA constructs, cells were treated with drugs 4-5 days after transduction; for overexpression constructs, cells were treated with drugs 3-5 days after transduction.

For both differentiated and undifferentiated cells, half of the medium was replaced every 2-3 days with fresh medium and cells were kept in an incubator under 7.5% CO2.

### Primary embryonic rat cortical neurons

Cortical neurons were grown on plates coated with poly-D-lysine (P1149, Millipore Sigma). 139 μL/cm^2^ of 0.1 mg/mL poly-D-lysine was added to multiwell plates, and plates were left covered in the cell culture incubator overnight. The following day, the poly-D-lysine solution was aspirated and the wells were washed at least three times with sterile deionized water. The wells were allowed to dry and cells were plated the same day.

Cortices were harvested from E17-E18 rat embryonic brains in cold HBSS without calcium chloride, magnesium chloride, or magnesium sulfate (Thermo Fisher Scientific) using a dissection microscope. The cortices were dissociated by a 10-minute incubation in 0.05% trypsin at 37°C and subsequent trituration in HBSS with two fire-polished pasteur pipettes of decreasing aperture size. Neurons were then passed through a cell strainer, counted, and plated at 5×10^5^ cells/mL on poly-D-lysine-coated plates in Neurobasal medium supplemented with 2% B27 supplement (Thermo Fisher Scientific), 500 mM Glutamax (Thermo Fisher Scientific), and 1% penicillin-streptomycin.

Half of the medium was replaced with fresh medium twice a week, and cells were kept in an incubator under 5% CO2. Cells were used for experiments after 6-9 days in vitro.

### SH-SY5Y cells

Human neuroblastoma SH-SY5Y cells were a kind gift from Dr. Ismael Santamaria Perez and were grown in DMEM/F12 medium (Cellgro) supplemented with 10% FBS, 2 mM glutamine, 1% penicillin-streptomycin under 5% CO2.

### Immortalized Mouse Embryonic Fibroblasts

Immortalized MEFs were a kind gift from Dr. Thong Ma from the Un Kang lab (Columbia University). The isolation and immortalization of these cells were described in [67]. Cells were grown in DMEM supplemented with 10% FBS and 1% penicillin-streptomycin under 5% CO2.

### Human Embryonic Kidney 293T cells

HEK293T cells were grown in DMEM supplemented with 10% FBS and 1% penicillin-streptomycin under 5% CO2. Cells were passaged every 3-4 days.

### Cloning

### shRNA constructs

Annealed oligomer sequences (IDT) for shRNAs were cloned into the pLVTHM vector. The latter was cut using MluI and ClaI before ligation with annealed oligomer. The shRNA target sequences used in this study were as follows: Control shRNA: 5’-GACCCTTGAATTGACTGTT-3’; Parkin: 5’-GGACACATCAGTAGCTTTG-3’; PINK1: 5’-T CAGGAGATCCAGGCAATT-3’.

### Tagged parkin under minimal parkin promoter

Two steps were involved in generating a “moderately overexpressed” parkin construct. First, rat parkin cDNA was amplified from a previously generated parkin construct (described in [14]) using primers to attach HA- and FLAG-tags on the N-terminus. The primers used were: F: ctagcctcgaggtttaaacATGTACCCATACGATGTTCCAGATTACGCTGCAGCAAATGATATCCTGGATTACAAGGATGACGACGATAAGctcccgcggacgcgtacgATAGTGTTTGTCAGG; R: tgcagcccgtagtttaaacctaCACGTCAAACCAGTGATC. This amplified, N-tagged cDNA was then inserted into a pWPI vector (Addgene) that had been linearized at the PmeI restriction site. In-Fusion cloning (Clontech) was used for this purpose, according to the manufacturer’s instructions.

In the second step, the EF-1α promoter of the pWPI-HA-FLAG-parkin construct was cut out at the PpuMI and PacI sites, and the minimal human parkin promoter amplified from a commercially available luciferase construct (SwitchGear Genomics) was inserted in its place using the NEBuilder^®^ HiFi DNA Assembly kit from New England Biolabs.

### Site-directed mutagenesis

Site-directed mutagenesis of “moderately overexpressed” parkin was carried out using the Q5 Site-Directed Mutagenesis kit from New England Biolabs, according to the manufacturer’s instructions.

### Lentiviral preparation

Early passage 293T cells were plated at a density of ~14-18 x 10^6^ cells/15cm plate. The next day, cells were transfected with 21 μg expression vector, 16.5 μg psPAX2, and 7.5 μg VSVg plasmids per plate using the calcium phosphate transfection method. Virus-containing medium was harvested 2 and 3 days after transfection, pooled, passed through a 0.45 μm filter, and concentrated using the Lenti-X concentrator (Clontech, #631231), according to the manufacturer’s instructions.

To evaluate viral titer, a range of viral solution volumes was added to differentiating PC12 cells, cells were fixed 3 days later, and GFP expression was evaluated by immunofluorescence. Viral infection efficiency was calculated by dividing the number of GFP-positive cells by the number of nuclei in the same field (assessed by Hoechst stain) using Fiji. For knockdown experiments, viral solution volumes yielding ≥70% infection efficiency were used, with an effort to match infection efficiencies between viruses. For exogenous parkin experiments, viral titer was assessed by Western blot, with viral volumes for subsequent experiments adjusted to achieve equal expression levels between constructs.

### L-DOPA, hydrogen peroxide, and CCCP treatments

Prior to drug treatment, half of the medium was removed from cells. An equal volume of fresh medium containing drugs at 2X the working concentration was added to cells to initiate the experiment.

For L-DOPA experiments, a 10 mM L-DOPA stock solution was freshly prepared before every experiment in 50 mM HCl and filter-sterilized. L-DOPA stock solution or an equal volume of 50 mM HCl was diluted in fresh medium before addition to cells. In experiments involving pretreatment of cells with an inhibitor, the inhibitor was added with fresh medium and L-DOPA was added directly to each well.

3% (w/v) hydrogen peroxide was diluted with sterile distilled water 1:1000 before subsequent dilution in fresh medium.

CCCP was dissolved in DMSO to yield a 10 mM stock solution, which was filter-sterilized and added to fresh medium before addition to cells. The stock solution was stored at - 80°C and used multiple times.

### Inhibitor & antioxidant treatments

Inhibitors and antioxidants were dissolved in water when possible and in DMSO when not water-soluble. In most cases, stock solutions of the drugs were diluted in fresh medium before addition to cells (with the exception of cycloheximide, in which case fresh medium was added to cells the day before treatment and cycloheximide stock solution was added directly to cells). In such cases, drugs and cellular stressors (L-DOPA, H2O2, CCCP) were added to cells at the same time. In the cases of carbidopa and cycloheximide, the latter were added to cells before treatment with L-DOPA. Cells were pretreated with carbidopa for 1.5 hours and cycloheximide for ~15 minutes before L-DOPA addition.

### Western blotting

Cell lysates were harvested in 1X Cell Lysis Buffer (Cell Signaling Technology) supplemented with Complete Mini protease inhibitor tablet –EDTA (Millipore Sigma) or directly in Western blot loading buffer (29% Cell Lysis Buffer, cOmplete Mini protease inhibitor tablet, 1X NuPage Reducing Agent (Thermo Fisher Scientific), 1X NuPage LDS Sample Buffer, 36% water). Lysis buffers were added directly to culture plate wells.

When harvested in Cell Lysis Buffer, lysates were kept on ice and subsequently sonicated with a probe sonicator at the lowest amplitude with a pattern of 1 second ON/ 1 second OFF, for a total of 10 seconds ON. The protein concentration in lysates was then determined using the Bradford assay using Protein Assay Dye Reagent Concentrate (Bio-Rad). Upon determining protein concentrations, samples were diluted with water to match the concentration of the most dilute sample and supplemented with NuPage Reducing Agent and NuPage LDS Sample Buffer. Following addition of Reducing Agent and LDS Buffer to cell lysates, these were boiled for 20 minutes in a dry heat bath at 95-100°C. After boiling, sample proteins were separated by SDS-PAGE and detected by Western blotting. The NuPAGE electrophoresis system was used, with precast Bis-Tris gels, MOPS SDS running buffer (or MES SDS running buffer for low molecular weight proteins), and NuPAGE transfer buffer. Generally, 4-12%gels were used and ~20 μg of protein was added per well. For one sample per gel, half the sample was added in a well adjacent to the full volume of sample to serve as a standard (see below). Proteins were transferred onto a nitrocellulose membrane for ~2 hours at 35V at 4°C. Following transfer, membranes were blocked for 1 hour at room temperature with gentle shaking in 5% milk in TBST (TBS + 1% Tween 20).

Membranes were probed with primary antibodies diluted in TBST with 5% BSA overnight at 4°C. The next day, membranes were subjected to 3 five-minute washes in TBST, incubated for 1 hour at room temperature with LI-COR IRDye secondary antibodies in 5% BSA/TBST in the dark, and washed three times again. Following incubation with secondary antibodies, membranes were kept in TBS in the dark.

Membranes were imaged using an Li-Cor Odyssey CLX scanner and band intensities were quantified using Image Studio Lite software (ver. 4.0.21). Briefly, after background subtraction, the signal from the lane in which a half volume of sample was loaded was assigned a value of “0.5” using the “concentration standard” feature. Similarly, the signal from the lane corresponding to full volume of sample was assigned a value of “1”. The program’s linear interpolation feature was then used to assign values to the remaining bands on the blot. This process was carried out both for proteins of interest and for loading controls, after with the former were normalized to the latter. For most experiments, bands of interest were normalized to the average of the corresponding ERK1 and ERK2 signals, though sometimes Ponceau stain was also used as a loading control.

### Subcellular fractionation

To prepare crude mitochondrial and cytosolic fractions from cells, the following protocol was followed. Cells and lysates were kept on ice for the duration of the procedure, and all centrifugation steps were performed at 4°C. First, cells were sprayed from the dish in PBS and pelleted by centrifugation. Cells from four wells of a 24-well plate were combined in one tube. 70 μL fractionation buffer was added to the cell pellet. The fractionation buffer consists of 220 mM mannitol, 70 mM sucrose, 20 mM Hepes-KOH (pH 7.5), 2 mg/ml BSA, 1 mM EDTA. Halt protease/phosphatase inhibitor (Thermo Fisher Scientific #78440) was added to the buffer right before use.

Cells were lysed by passage through a 27G needle thirty times, until there were about 8 broken nuclei for every unbroken cell. To remove cell debris & nuclei, samples were centrifuged at 1,000 x g for 10 minutes. The supernatant was set aside and the nuclear fraction was washed with 60 μL fractionation buffer and centrifuged a second time. The supernatant from this second spin was combined with the supernatant from the first spin and centrifuged at 10,000 x g for 10 minutes to separate the mitochondria-rich pellet from the cytosolic fraction. The mitochondria-rich pellet was washed three times by resuspending the pellet in 60 μL fractionation buffer and re-pelleting at 10,000 x g for 10 minutes. Then, the mitochondrial pellet was resuspended in 13 μL Cell Lysis Buffer (Cell Signaling Technology) with cOmplete Mini Protease inhibitor (Millipore Sigma). The mitochondrial and cytosolic fractions were supplemented with NuPAGE Reducing Agent and LDS sample buffer (Thermo Fisher Scientific), boiled for 20 minutes at 95°C, and used for Western blotting following the standard protocol.

### qPCR

Analysis of mRNA by qPCR was performed as in [14]. Briefly, total cellular RNA was extracted using TRI reagent (Molecular Research Center). cDNA was synthesized using the first-strand cDNA synthesis kit (Origene). qPCR was performed using FastStart SYBR Green Master Mix (Roche) and an Eppendorf Realplex Mastercyler. The following cycling protocol was used: 1 cycle at 95°C for 10 min and 40–45 cycles of amplification: 95°C for 15 s, 58–60°C for 30–60 s, 72°C for 30–60 s. Relative mRNA amounts were determined using the delta-delta Ct method with 18S rRNA as the housekeeping gene. The following primers were used: Parkin: F: GGCCTTTGCAGTAGACAAAA; R: ACCACAGAGGAAAAGTCACG; 18S: F: TTGATTAAGTCCCTGCCCTTTGT; R: CGATCCGAGGGCCTCACTA; PINK1: F: AAAGGCCCAGATGTCGTCTC; R: GCTTAAGATGGCTTCGCTGG.

### High Performance Liquid Chromatography

Cells were washed twice with cold PBS, detached from wells via forceful pipetting, and pelleted at maximum speed in a table-top centrifuge for 5 minutes at 4°C. The supernatant was then removed and the cell pellet was resuspended in 75 μL PBS. 25 μL of this cell mixture was removed, pelleted, and resuspended in 40 μL Cell Lysis Buffer with protease inhibitors. These samples were then sonicated, and the Bradford assay was performed to determine protein concentrations. The remaining 50 μL of cell mixture was mixed with an equal volume of 0.5 M trichloroacetic acid and vortexed for 10 seconds. These samples were then centrifuged at 10,000 x g for 2 minutes at 4°C, and the supernatant was transferred to a new tube. This supernatant was used for HPLC.

Samples were separated on Brownlee VeloSep RP-18, 3 μm, 100 x 3.2 mm column (Perkin Elmer, Waltham, MA) using Gilson (Middleton, WI) isocratic 307 pump. The mobile phase consisted of 45 mM NH_2_PO_4_, pH 3.2, 0.2 mM EDTA, 1.4 mM heptanesulfonic acid and 5% methanol. L-DOPA peaks were detected using an ESA (Chelmsford, MA) Coulochem II electrochemical detector at 300 mV oxidation potential. Relative L-DOPA levels were calculated by normalizing the area under the L-DOPA HPLC peak to the protein concentration from the same sample.

### Immunofluorescence (for quantification of viral titer)

Cells were fixed for 15 min in 4% paraformaldehyde and then washed 3 times with 1X PBS. Cells were then blocked with Superblock blocking buffer supplemented with 0.3% Triton X-100 for 1-2 h at room temperature and incubated overnight at 4°C with primary antibody in Superblock/Triton X (chicken anti-GFP, 1:1000, #A10262, Thermo Fisher Scientific) with gentle shaking. The next day, primary antibody was washed off in 3×8-minute washes with gentle shaking using 1X PBS. Cells were then incubated with fluorescent secondary antibody in Superblock/Triton = for 1 hour with gentle shaking (AlexaFluor-488 anti-chicken, 1:1000, #A11039, Thermo Fisher Scientific). Cells were then washed 3 = 8 minutes in 1X PBS with gentle shaking. Hoechst 33328 was added in the first wash (1:2500). Cells were kept in 1X PBS before imaging using an inverted fluorescence microscope.

### Quantification of cell death

Cell death was assessed as previously described [129]. Briefly, cells were treated with a lysis buffer (150 μL lysis buffer per cm^2^) that disintegrates the plasma membrane while keeping nuclei intact and counting viable nuclei manually using a hemacytometer. At least 200 nuclei were counted per condition. 10X lysis buffer recipe: 5 g of cetyldimethyl-ethanolammonium bromide, 0.165 g of NaCl, 2.8 ml of glacial acetic acid, 50 ml of 10% Triton X-100, 2 ml of 1 M MgCh, 10 ml of 10X PBS, and 35.2 ml of H_2_O.

### Statistical analysis

Statistical analysis was performed using GraphPad Prism 6 and using an online multiple comparison correction resource (http://alexandercoppock.com/statistical_comparisons.html) developed by Alexander Coppock, as described in [130].

For most experiments, values were normalized to the control condition in that experiment before replicate experiments were combined. Paired t-tests were performed in most cases, followed by Holm correction for multiple comparisons. In time-course analyses, unpaired t-tests at each time point followed by Holm-Sidak’s multiple comparisons test were used. For analysis of differences between parkin mutants, one-way ANOVA of stressor-treated mutants was performed, followed by Holm-Sidak correction for multiple comparisons. For analysis of the cycloheximide experiment, repeated measures 2-way ANOVA of data from the 12-48 hour time points was used with Sidak’s multiple comparisons test.

## Acknowledgements

MEFs were a gift from the Un Kang lab (Columbia University) and SH-SY5Y cells were a gift from Ismael Santa-Maria Perez (Columbia University). We thank Carlo Corona from Michael Shelanski’s lab (Columbia University) and Chandler Walker from Ulrich Hengst’s lab (Columbia University) for assistance with primary cortical neuron cultures. This work was supported by the NIH.

---

**Supplementary Figure 1.**
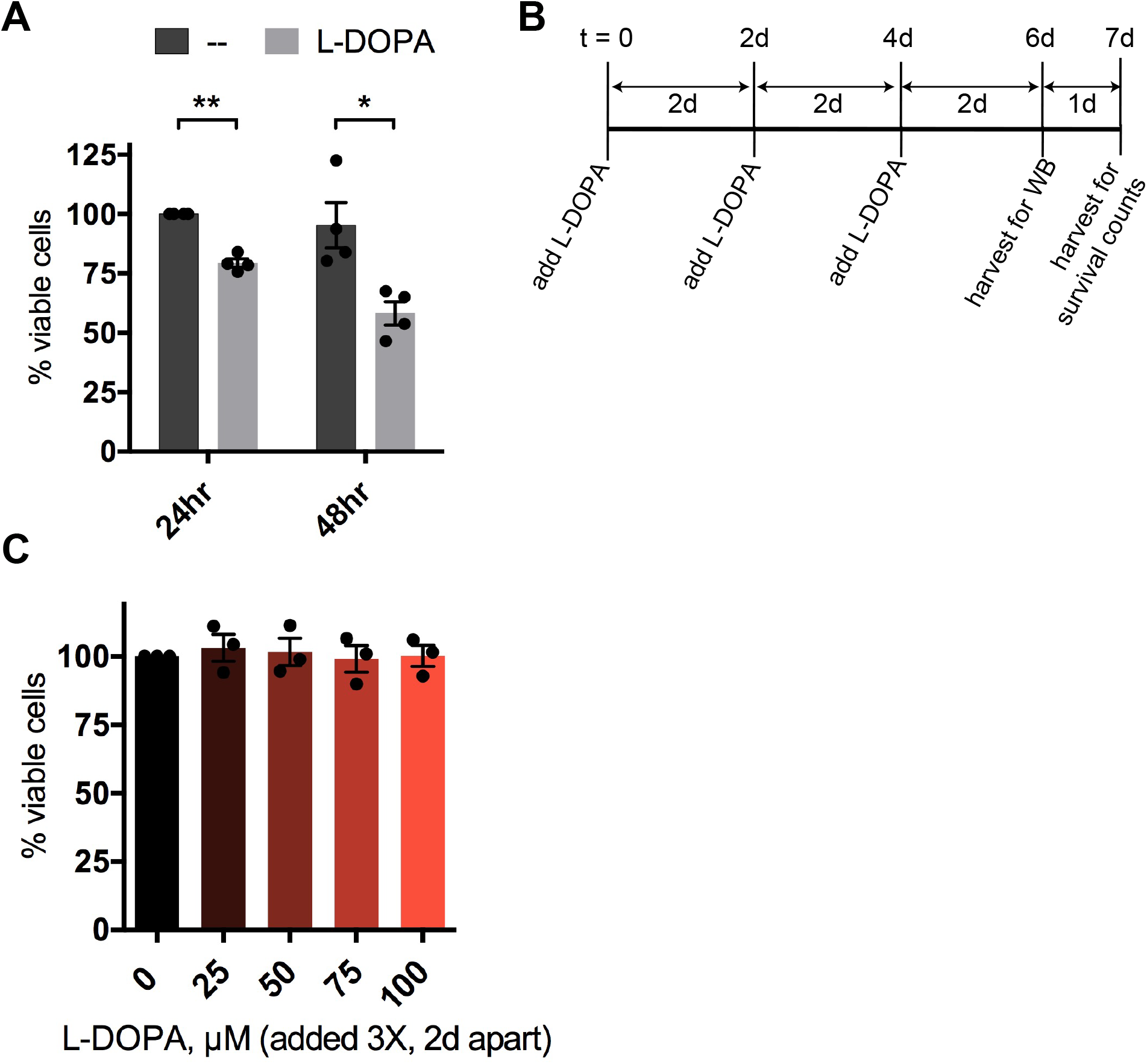
L-DOPA-induced parkin protein loss occurs independently of cell death. A. Timecourse of L-DOPA toxicity in differentiated PC12 cells. Cells were treated with 200 μM L-DOPA and harvested for assessment of survival after 24 and 48 hours. B. Treatment paradigm used to assess whether non-toxic doses of L-DOPA lead to parkin loss. WB=Western immunoblot. C. Quantification of cell survival after exposure to L-DOPA following the paradigm in (B). A,C. Error bars show SEM from N = 4 (A) or 3 (C) independent experiments; * p ≤ 0.05, ** p ≤ 0.01 by paired t-test with Holm correction for multiple comparisons.

**Supplementary Figure 2.**
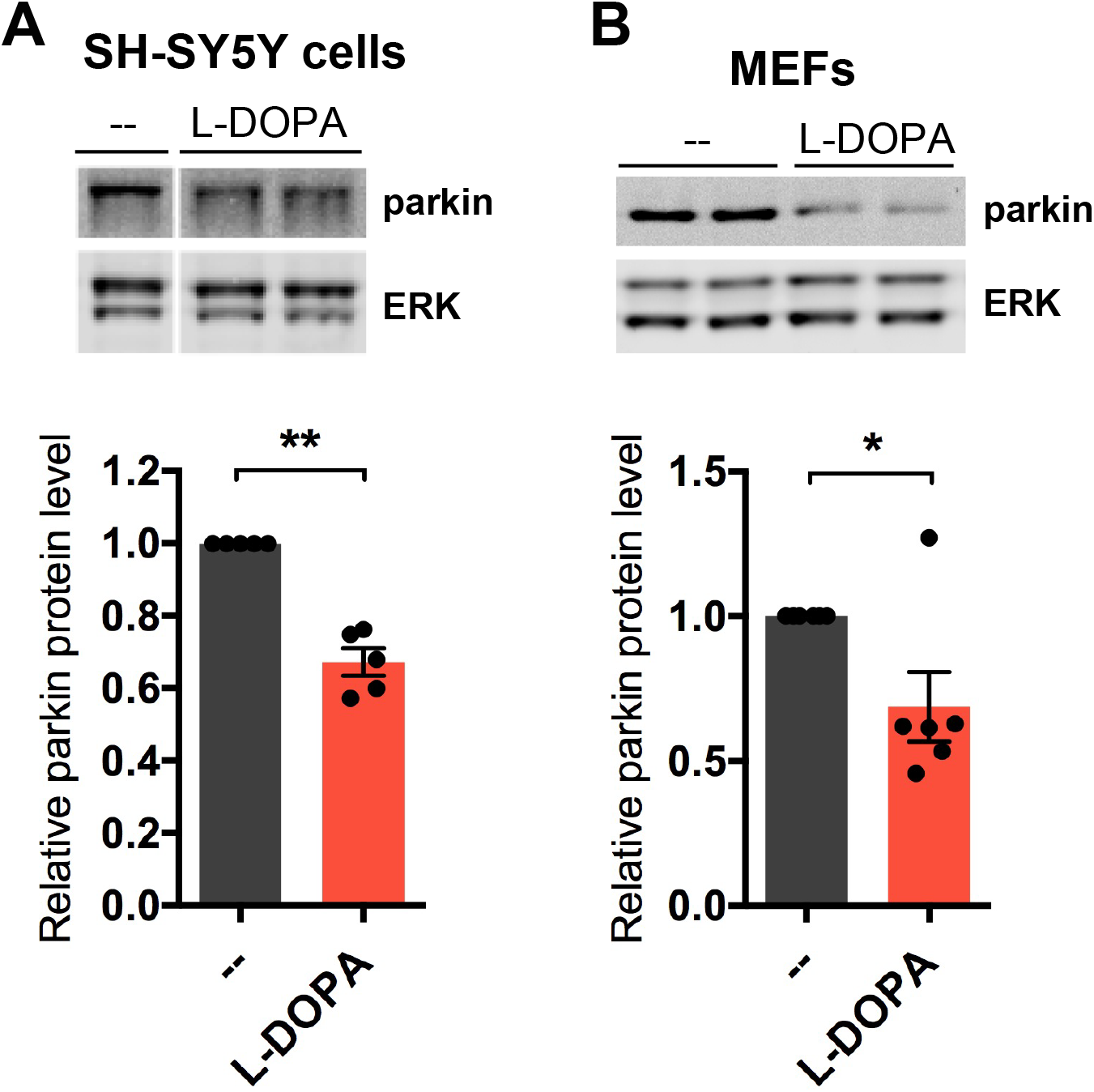
L-DOPA treatment induces parkin protein loss in SH-SY5Y cells and mouseembryonic fibroblasts. L-DOPA induces parkin loss in undifferentiated SH-SY5Y cells (A) and immortalized MEFs (B). Cells were treated with 200 μM L-DOPA for 24 hours before being harvested for Western immunoblotting. Error bars show SEM from N = 5 (A) or 6 (B) independent experiments; * p ≤ 0.05 by paired t-test.

**Supplementary Figure 3.**
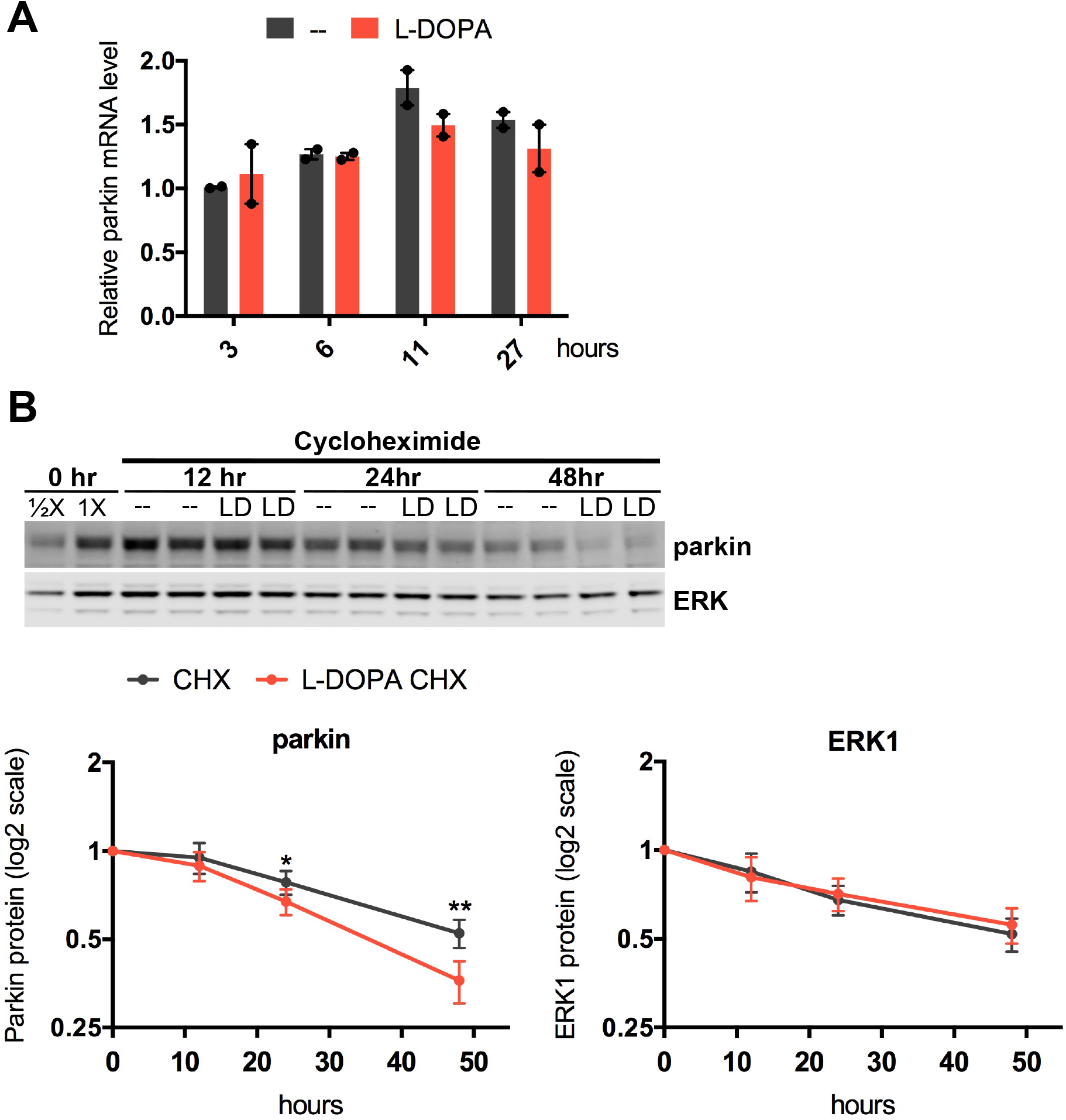
L-DOPA treatment increases the rate of parkin protein turnover. A. L-DOPA does not alter parkin mRNA levels. Cells were treated with 200 μM L-DOPA and harvested for qPCR analysis at the indicated time points. The parkin mRNA levels were normalized to 18S ribosomal RNA. B. L-DOPA accelerates parkin loss in PC12 cells when translation is inhibited. Cells were treated with 2 μg/mL cycloheximide (CHX) in the presence or absence of 200 μM L-DOPA (LD) and harvested for Western blotting at the indicated time points. Representative Western blot and quantification of parkin and ERK1 levels not normalized to a loading control. Error bars show range from N = 2 independent experiments for (A) and SEM from N = 3 independent experiments for B; For A, significance was assessed at each time point by paired t-test followed by Holm correction for multiple comparisons. For B, * p ≤ 0.05, ** p ≤ 0.01 relative to no L-DOPA at that time point by repeated measures 2-way ANOVA of data from the 12-48 hour time points with Sidak’s multiple comparisons test.

**Supplementary Figure 4.**
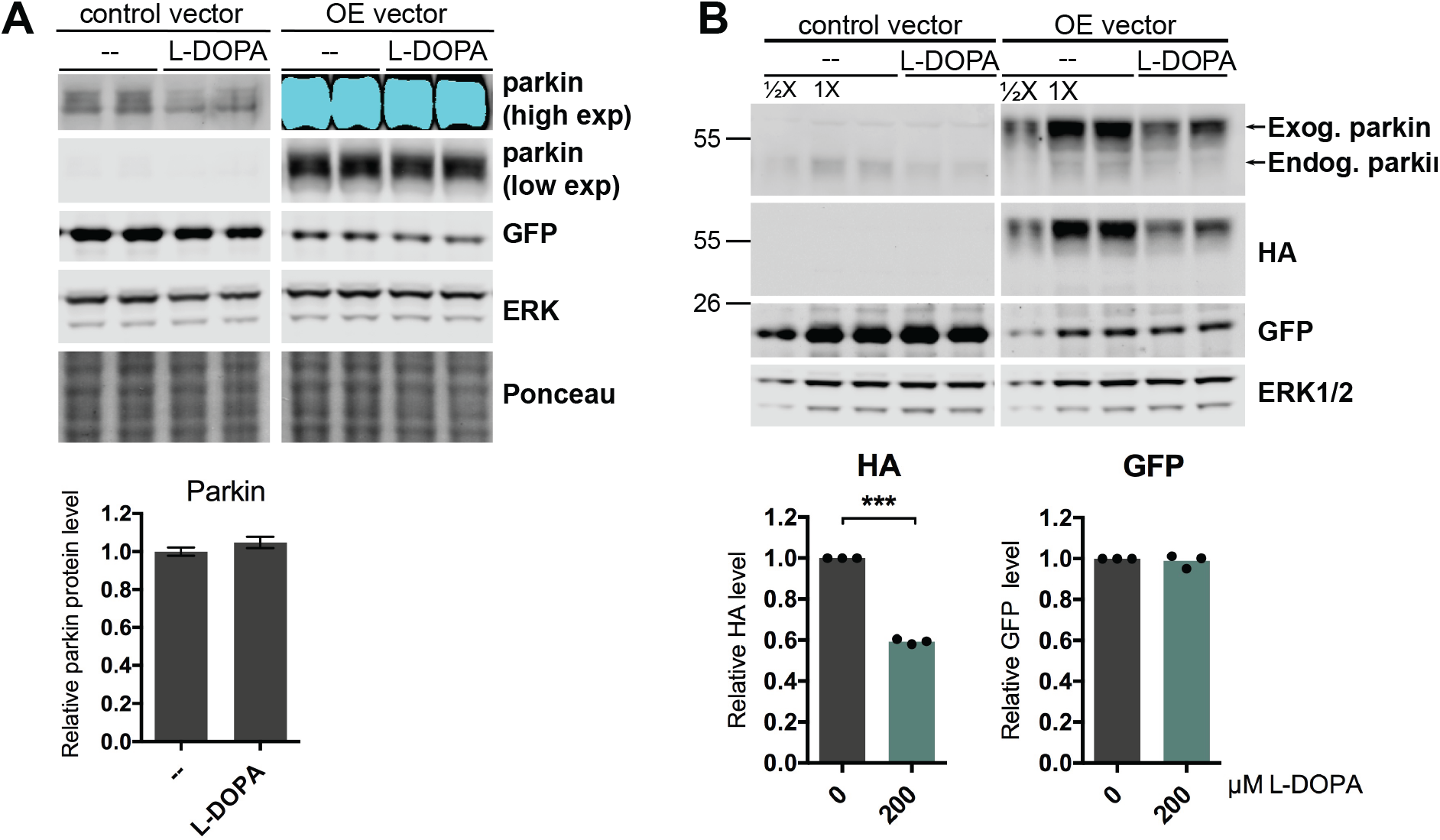
L-DOPA treatment decreases moderately overexpressed parkin protein. A. Differentiated PC12 cells were transduced with a lentiviral vector carrying untagged rat parkin expressed from the EF-1α promoter (OE vector). GFP is expressed bicistronically with parkin using an IRES sequence. 16 hours after transduction, cells were treated with 200 μM L-DOPA and harvested for WB 48 hours later. Parkin protein levels were quantified relative to the average of ERK1 and Ponceau stain signals. Error bars in B show the range of the two biological replicates shown in the immunoblot. Results are representative of 2 experiments. B. Differentiated PC12 cells were transduced with a lentiviral vector carrying “moderately” overexpressed, N-terminally HA- and FLAG-tagged rat parkin expressed from the minimal human parkin promoter (OE vector). GFP is expressed bicistronically with parkin using an IRES sequence. 3 days after transduction, cells were treated with 200 μM L-DOPA for 24 hours and harvested for WB. Overexpressed parkin protein levels were detected using antibodies against parkin and HA. Quantification of overexpressed parkin and GFP levels with and without 200 μM L-DOPA treatment from N = 3 independent experiments is shown. Proteins were normalized to the average of ERK1 and 2. Error bars showing SEM are too small to be seen. *** p ≤ 0.001 by paired t-test. A,B. Control vector was an empty version of the parkin construct used in A.

**Supplementary Figure 5.**
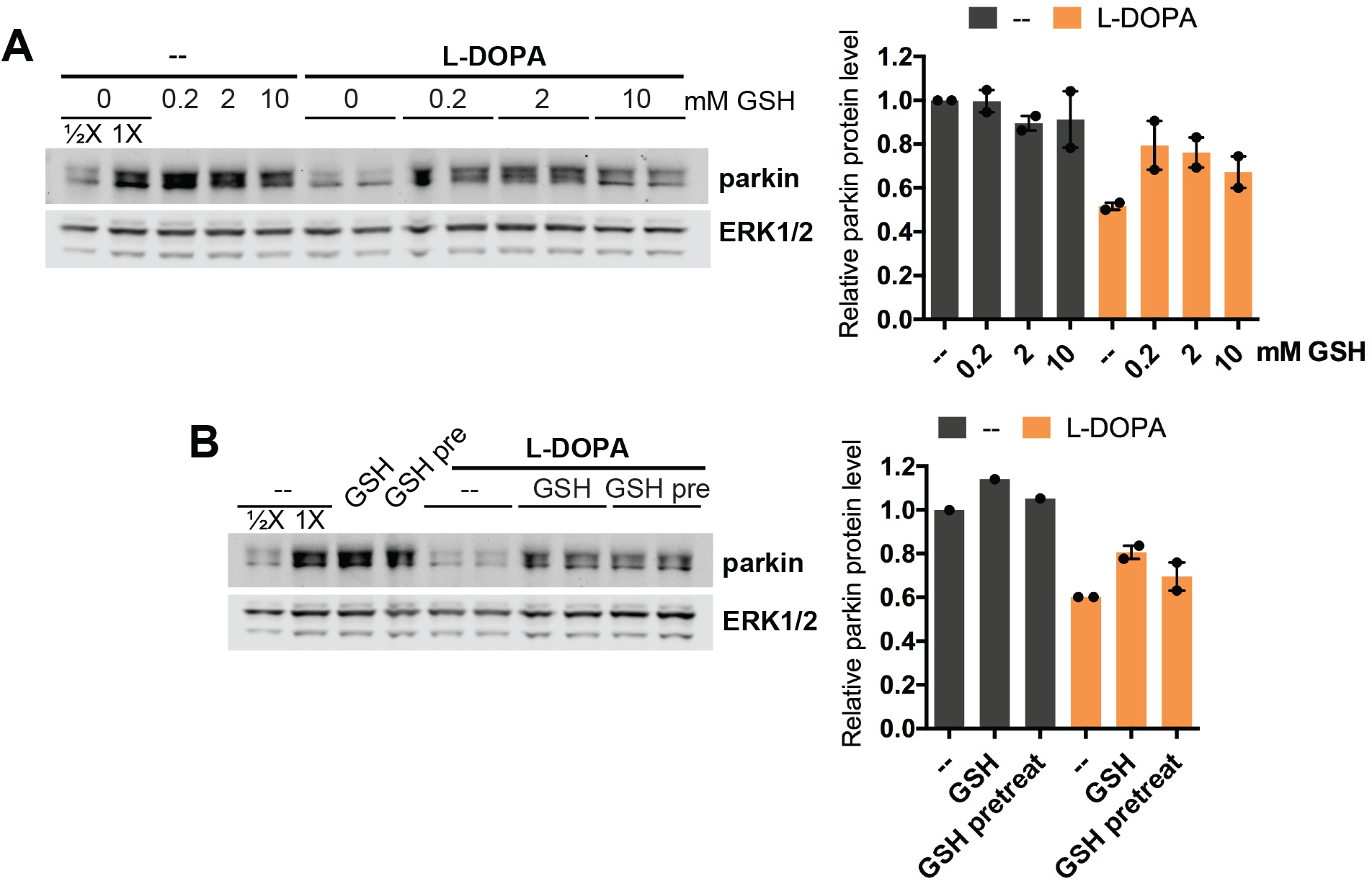
Effect of high glutathione doses and glutathione pre-treatment on L-DOPA-induced parkin loss. A. Differentiated PC12 cells were co-treated with 200 μM L-DOPA and the indicated glutathione (GSH) concentrations for 24 hours before lysates were harvested for Western blotting. A representative Western blot and quantification of parkin levels from 2 independent experiments are shown. B. PC12 cells were co-treated with 200 μM L-DOPA and 200 μM GSH or pre-treated with 200 μM GSH for 4 hours prior to addition of L-DOPA (“GSH pre”). Data points in the quantification of parkin levels represent biological replicates from one experiment. Error bars show range.

**Supplementary Figure 6.**
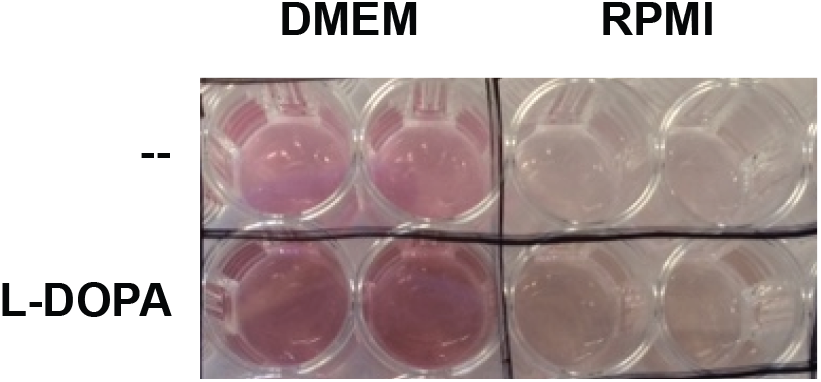
L-DOPA causes browning of cell culture medium. 200 μM L-DOPA was added to DMEM or RPMI culture medium and incubated for 24 hours at 37°C under 7.5% CO2 in the absence of cells. Evidence of L-DOPA-derived quinones can be seen in the “browning” of the medium.

**Supplementary Figure 7.**
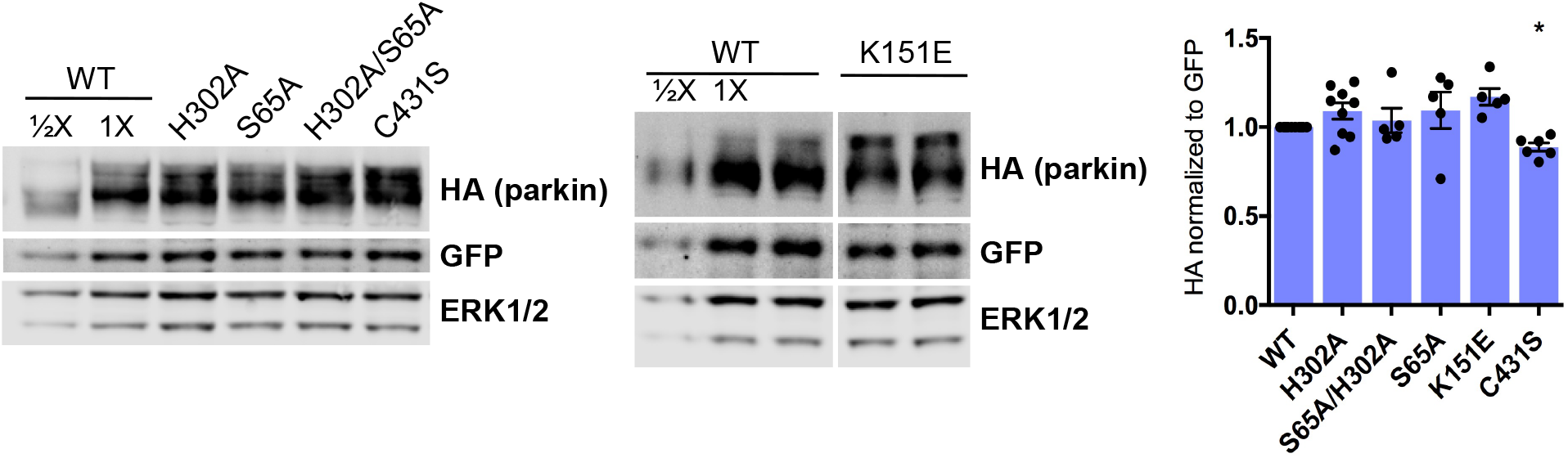
Expression levels of parkin mutants are similar to one another. Differentiated PC12 cells were transduced for 3 to 6 days with lentiviral vectors carrying the indicated parkin mutants and bicistronically expressed GFP before cells were harvested for Western immunoblotting. Representative blots are shown. To compare relative expression levels of the different parkin mutants, parkin levels for each construct were normalized first to ERK1 and 2 to control for loading differences and then to GFP to control for variability in viral titer. The results of this quantification are shown on the right. Error bars show SEM from 5-10 independent experiments; * p ≤ 0.05 relative to WT by paired t-test with Holm correction for multiple comparisons.

**Supplementary Figure 8.**
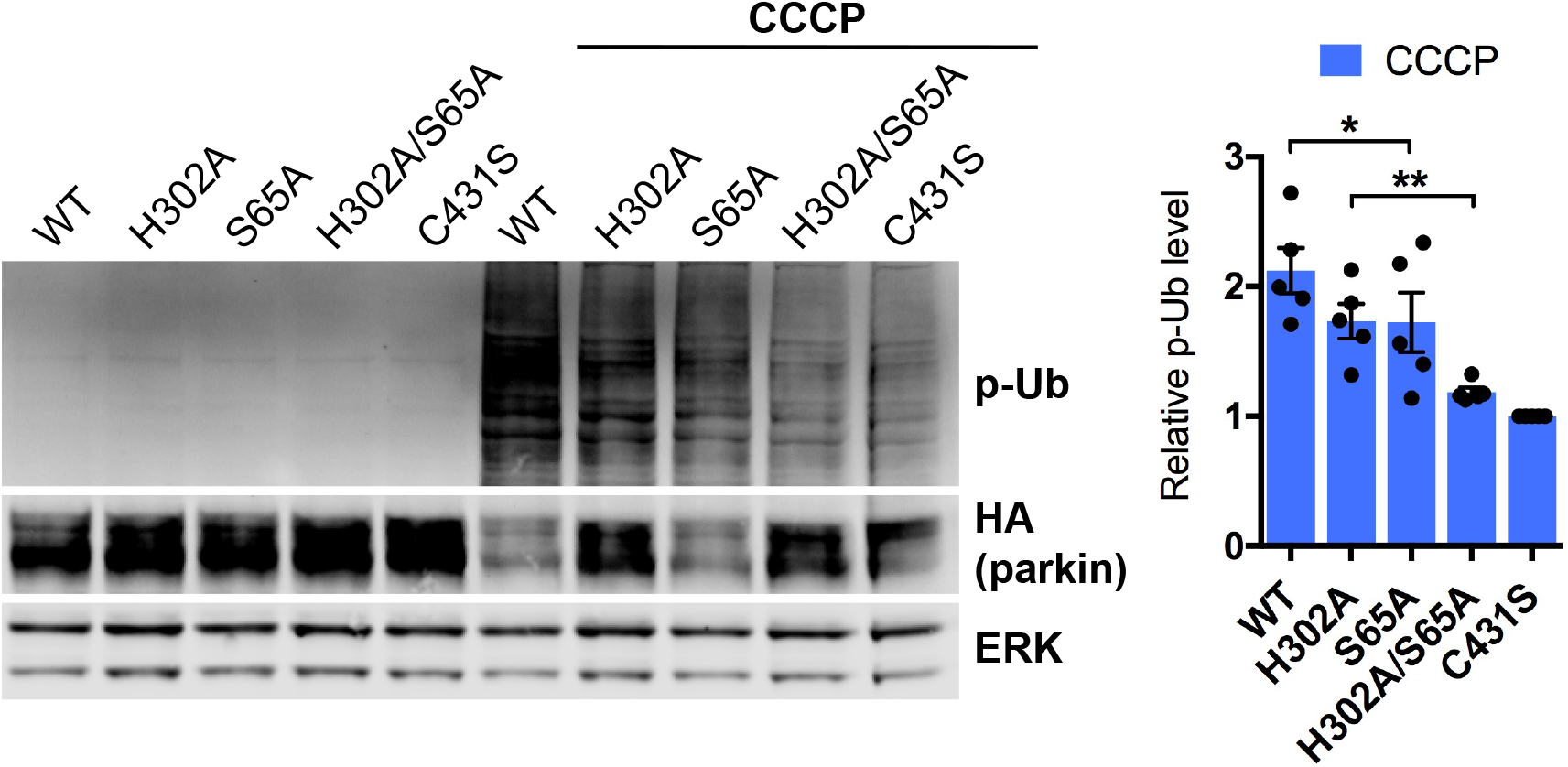
Parkin phosphorylation plays a functional role in the CCCP model. Differentiated PC12 cells were transduced with indicated mutant parkin constructs and treated with CCCP for 6 hours as described in Fig. 5. Relative levels of phospho-poly-Ub for each mutant were quantified by normalizing the p-Ub signal first to ERK1/2 and then to the corresponding mutant parkin level in untreated cells as determined by probing with anti-HA (to control for infection efficiency). C431S is catalytically inactive parkin. Quantification from N = 5 independent experiments is shown. Error bars show SEM. * p ≤ 0.05, ** p ≤ 0.01 by paired t-test with Holm correction for multiple comparisons.

**Supplementary Figure 9.**
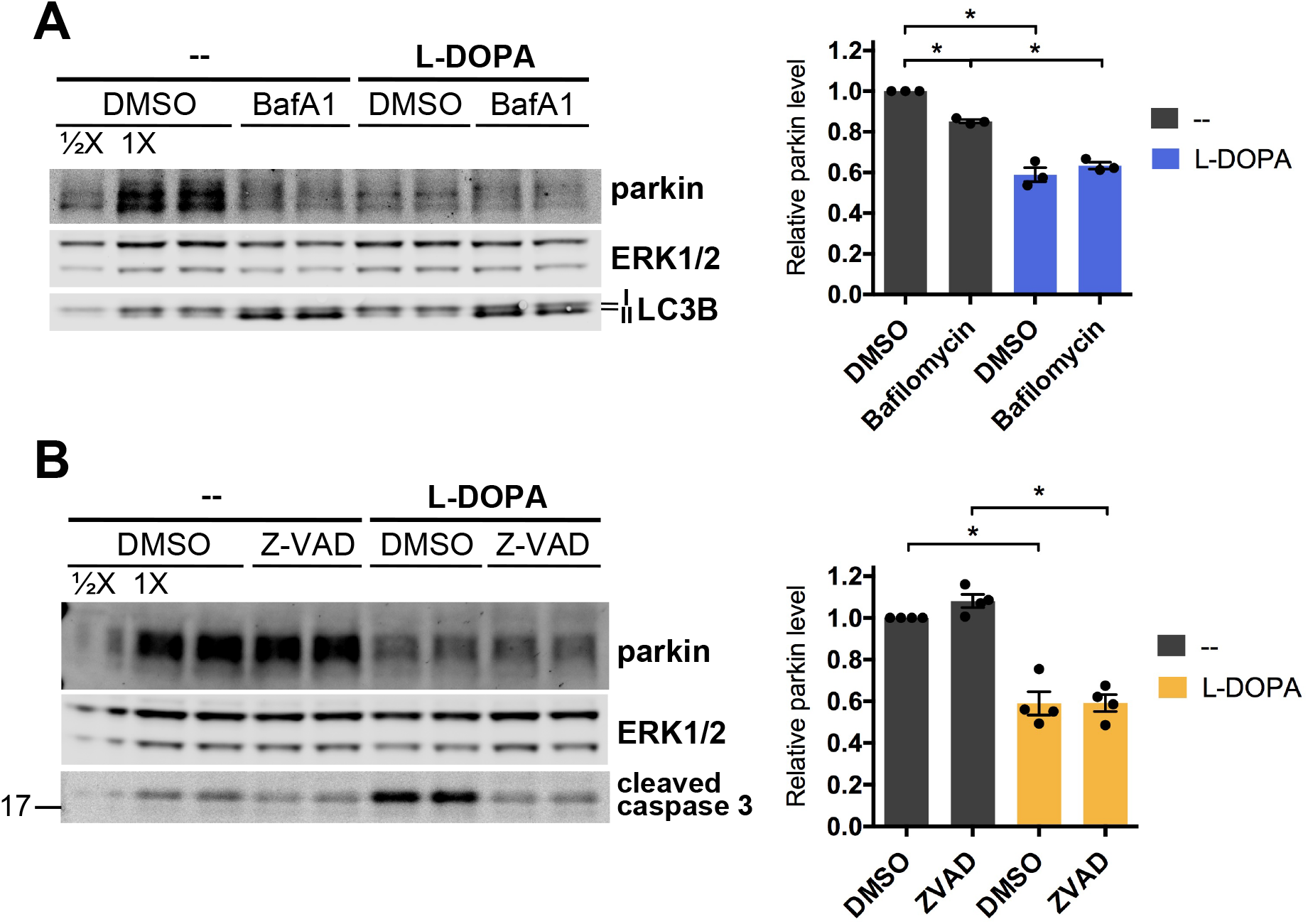
L-DOPA-induced parkin loss is not attenuated by lysosomal inhibition orcaspase inhibition. A. Inhibition of lysosomal degradation doesn’t prevent parkin protein loss from L-DOPA treatment. Differentiated PC12 cells were co-treated with 200 μM L-DOPA and 50 nM bafilomycin A1 (BafA1) for 24 hours before harvest for Western immunoblotting. Increased LC3B II signal was a positive control for the effectiveness of bafilomycin A1. B. Inhibition of caspase cleavage doesn’t prevent parkin loss from L-DOPA treatment. Differentiated PC12 cells were co-treated with 200 μM L-DOPA and 100 μM Z-VAD-FMK (Z-VAD) for 24 hours before cells were harvested for Western immunoblotting. Prevention of the formation of cleaved caspase 3 was a positive control for the effectiveness of Z-VAD-FMK. A,B. Representative blots and quantifications of N = 3 (A) and 4 (B) independent experiments are shown. Error bars show SEM; * p ≤ 0.05 by paired t-test with Holm correction for multiple comparisons.

**Supplementary Figure 10.**
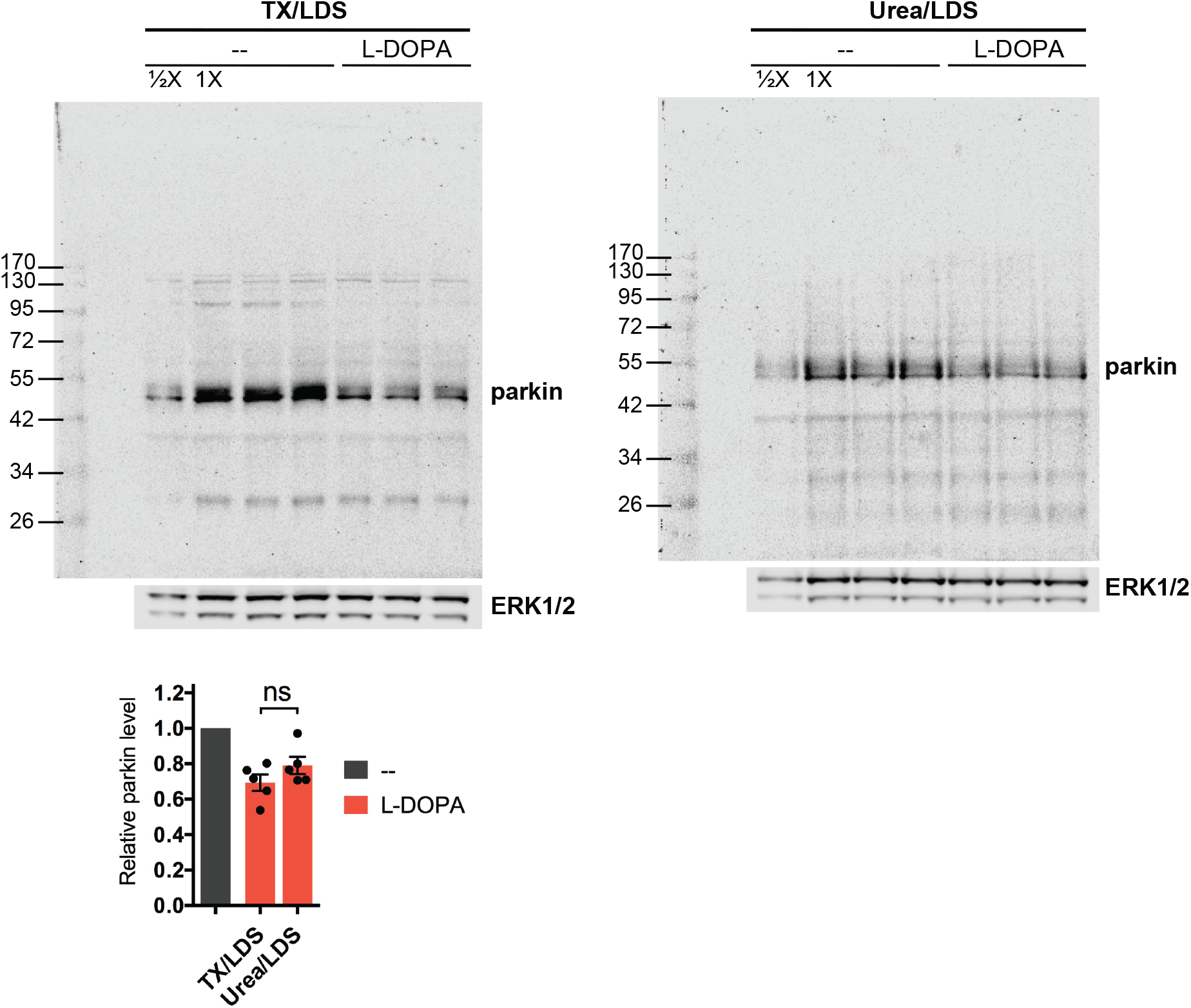
L-DOPA-induced parkin loss is not attenuated by harsher cell lysis. Parkin is lost to the same extent after L-DOPA exposure whether cells are harvested in buffer containing Triton X-100 (TX) and 2% LDS (left immunoblot) or 8 M urea and 2% LDS (right immunoblot). Differentiated (N = 3) or undifferentiated (N = 2) PC12 cells were treated with 200 (N = 4) or 250 (N = 1) μM L-DOPA for 24 hours before being harvested in buffer containing Triton X-100 or 8 M urea, sonicated, supplemented with LDS and DTT, and subjected to Western immunoblotting. Parkin levels in L-DOPA-treated samples for each buffer were normalized to parkin levels in control samples in the same buffers; the gray bar in the quantification represents parkin levels in the control samples for each buffer. Error bars show SEM; statistical significance was queried by paired t-test with Holm correction for multiple comparisons.

